# Tau deletion results in sex-dependent modulation of synaptic weakening in rat hippocampus

**DOI:** 10.1101/2025.08.07.665155

**Authors:** Liam T. Ralph, Patrick Tidball, Eric W. Salter, Brian Park, Lijia Zhang, Mengdi Wei, Lauren A. Joe, Jonathan S. Thacker, Sun-Lim Choi, Ashish Kadia, Fuzi Jin, Meng Tian, Clarrisa A. Bradley, Paul E. Fraser, Zhengping Jia, Gerold Schmitt-Ulms, Marina Gertsenstein, Lauryl M.J. Nutter, John Georgiou, Graham L. Collingridge

## Abstract

Tangles, a defining characteristic of Alzheimer’s disease (**AD**), are composed principally of hyperphosphorylated and misfolded tau species. However, dysregulation of tau precedes tangle formation by many years and is associated with cognitive decline, the best functional correlate of which is synaptic weakening and eventual synaptic loss. Although tau is present at synapses, its normal synaptic function is poorly understood – information vital to the rational development of tau-modifying therapies. Since rats have a greater cognitive repertoire than mice and display more robust signs of tau pathology in AD models, we developed a tau knockout (***Mapt^-/-^***) rat to investigate the impact of tau elimination on synaptic function. We observed that long-term depression (**LTD**), a form of synaptic weakening, is enhanced in the hippocampus of male, but not female, rats. This enhanced LTD was dependent on the synaptic activation of group I metabotropic glutamate receptors and involved altered synaptic actin polymerization. These studies therefore provide mechanistic insights into how tau modulates synaptic function via regulation of the actin cytoskeleton. Our findings are relevant to the understanding of the physiological roles of synaptic tau, the functional consequences of tau-lowering therapies, and the influence of sex in AD susceptibility.

## INTRODUCTION

Tau, a microtubule-associated protein crucial for the maintenance of neuronal cytoskeletal architecture, is disrupted in tauopathies such as Alzheimer’s disease (**AD**)^1–4^. Beyond the canonical pool of tau in the axonal compartment, tau is also translocated to^5–7^ and present at^8,9^ synapses where it regulates synaptic plasticity under normal physiological conditions^10–13^. Since synaptic weakening and loss of synapses are early features of tauopathies^14,15^, deciphering the physiological role of tau in long-term depression (**LTD**), a form of activity-dependent synaptic weakening^16,17^, is imperative for understanding the aetiology of, and developing more efficacious therapeutics for, neurodegenerative diseases^18^. To date, the impact of tau deletion on synaptic function has primarily been investigated in microtubule-associated protein tau homozygous knockout (***Mapt^-^*^/-^**) mice, where tau deletion results in reductions in both LTD^8^ and long-term potentiation (**LTP**)^12,13^, processes that are necessary for learning, memory and other cognitive functions^19^.

Owing to the growing interest in rat models for investigating neurodegenerative diseases^20,21^, we generated and characterised a *Mapt^-^*^/-^ rat model to understand the physiological function of tau protein. To our surprise, we observed enhanced low-frequency stimulation (**LFS**)-induced LTD in male, but not female, juvenile *Mapt^-^*^/-^ rats. In contrast, we found there was reduced LTD in male *Mapt^-^*^/-^ mice in line with a previous report^8^, highlighting a species difference in the impact of tau deletion on synaptic function. We found that the enhanced component of LTD in male *Mapt^-^*^/-^ rats was mediated by group I metabotropic glutamate receptors (**mGluR_I_s**) and was independent of *N*-methyl-D-aspartate receptors (**NMDARs**). Furthermore, *Mapt^-^*^/-^ rats exhibited increased synaptic levels of actin, and inhibition of actin polymerization in male wildtype (**WT**) rats phenocopied the enhanced LTD in male *Mapt^-^*^/-^ rats. Together, these findings indicate that tau deletion in rats results in a sex-specific enhancement of LTD through an interplay between synaptic mGluR_I_ signaling and the actin cytoskeleton. Our results not only provide mechanistic insights into the sex-dependent physiological function of tau at synapses, but also have implications for the aetiology of tauopathies and the design of tau-lowering therapies.

## RESULTS

### Creation of a *Mapt^-/-^* rat model

To study the impact of tau deletion in rats, we developed a constitutive Crl:CD Sprague Dawley *Mapt^-^*^/-^ knockout rat line (**SD-*Mapt*^em1Tcp^**) using CRISPR-Cas9 (Fig. 1a), as per methodology described in^22^. Sanger sequencing confirmed deletion of the fourth and fifth exons of the *Mapt* gene based on the NCBI reference sequence for full-length *Rattus norvegicus* CNS tau (XM_008768277.3; Extended Data Fig. 1a). To confirm if this was sufficient to eliminate the translation of tau proteins, levels of tau were assayed in whole brain homogenates from juvenile (postnatal day (**P**) 14-19) rats by immunoblotting with an *N*-terminal tau monoclonal antibody. Accordingly, tau protein expression was absent in *Mapt^-^*^/-^ rats and reduced by approximately 50% in heterozygous (*Mapt^+/-^*) rats relative to WT rats (Extended Data Fig. 1b). Furthermore, tau was present in synaptically-enriched hippocampal fractions from juvenile WT rats, and its deletion was not associated with changes in levels of microtubule-associated protein (**MAP**) 1A, 1B, 2B, or 2C in either total or synaptically-enriched hippocampal fractions (Fig. 1b,c and Extended Data Fig. 2). We did not observe any gross defects in *Mapt^-^*^/-^ or *Mapt^+/-^* rat pup survival or development. Pups were born at expected Mendelian ratios from *Mapt^+/-^* breeding pairs (Fig. 1d), and body, brain and hippocampal mass were unaltered in *Mapt^-^*^/-^ rats (Fig. 1e). In summary, we have established a viable tau knockout rat model that does not impact juvenile expression of other MAPs.

**Figure 1.**
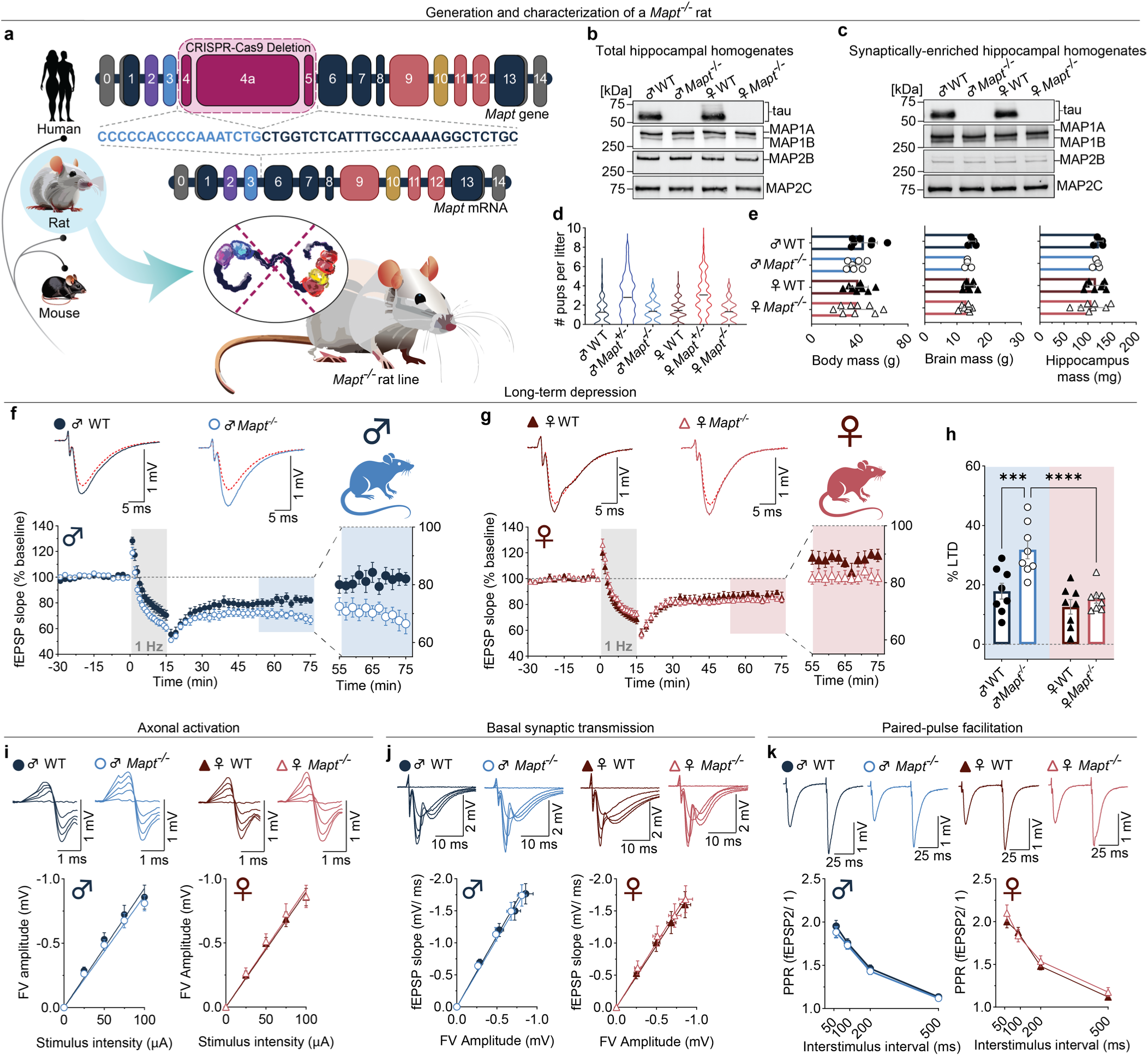
Male-specific enhancement of LTD in a novel *Mapt*^-/-^ rat. **a**, Schematic depicting the selection of animal models and deleted region in the rat *Mapt* gene of the SD-*Mapt*^em1Tcp^ rat line. **b**,**c**, Representative immunoblots showing levels of tau, MAP1A, MAP1B, MAP2B and MAP2C in total (**b**) and synaptically enriched (**c**) hippocampal fractions from male and female WT and *Mapt^-/-^* rats. The detection of tau in synaptically enriched fractions from WT rats suggests it is present at synapses of juvenile rats. Tau was absent in both total homogenates and synaptically enriched fractions from *Mapt^-/-^* rats, and tau deletion did not impact expression levels of other MAPs (*n* = 8 in all groups; quantification is shown in Extended Data Fig. 2). **d**, Both sexes of WT, *Mapt^+/-^* and *Mapt^-/-^* rats were born at expected Mendelian ratios (0.9:2.1:1.0 and 1.0:2.1:0.9 for males and females respectively, in litter sizes of 11.3 ± 0.4 rats; *n* = 144 litters). **e**, Absence of tau in both sexes of juvenile rats did not impact body, brain, or hippocampus mass (left-to-right, respectively; *n* = 4-7). **f-h**, Field potential recordings showing LTD induced by low frequency stimulation (LFS; 900 pulses delivered at 1 Hz) in the CA1 region of hippocampal slices prepared from male (**f**; circles) and female (**g**; triangles) WT (closed symbols) and *Mapt^-/-^* (open symbols) littermate pairs of juvenile rats. Expanded regions of the time plots show the final 20 min of recordings. Tau deletion had a sex-dependent impact on LTD (**h**; two-way ANOVA, genotype × sex interaction: *F*_(1, 28)_ = 5.09, *P* = 0.03; *n* = 8 in all groups). Specifically, male *Mapt^-/-^*rats exhibited enhanced LTD relative to male WT rats (Šídák’s post hoc test: *P* = 0.001), while LTD in female *Mapt^-/-^* rats was similar to that in female WT rats (*P* = 0.8). There were no significant differences in LTD between sexes of WT rats (*P* = 0.9). Representative traces show superimposed fEPSPs from the baseline period (solid line) and the end of the recording session (dashed line). The level of LTD was calculated as the percent change from baseline in the last 6 min of recordings. **i-k**, Axon connectivity and basal synaptic properties in the hippocampal CA1 region were unchanged in both male and female juvenile *Mapt^-/-^* rats. The plots show the input/output (I/O) relationship between the FV amplitude and stimulus intensity (**i**), the I/O relationship between the fEPSP slope and FV amplitude (**j**) and paired-pulse facilitation (PPF) measured across a range of interstimulus intervals (**k**) (*n* = 10 in all groups; quantification is shown in Extended Data Fig. 3). Traces above the I/O plots show superimposed FVs or fEPSPs elicited over a range of stimulus intensities. Traces above the PPF plots show fEPSP pairs evoked with a 50 ms interstimulus interval. Detailed statistics are shown in Supplementary Tables 1-3. In this, and subsequent figures, pooled data are presented as the mean ± s.e.m. \*\**P* < 0.01.

### *Mapt^-/-^* rats exhibit a sex-specific enhancement of LTD

Synaptic weakening is relevant to neurodegenerative diseases involving tau^14,23–25^, and previous work has shown inhibition of hippocampal NMDAR-dependent LTD in *Mapt* knockout mice^8^. We therefore assessed LTD of synaptic transmission induced by LFS (900 pulses delivered at 1 Hz) at CA3–CA1 synapses of hippocampal slices prepared from juvenile male and female *Mapt^-^*^/-^ rats and their WT littermate counterparts. Surprisingly, and in contrast to mice, we found that the absence of tau in rats resulted in an enhancement of LTD that was dependent on sex (two-way ANOVA: main effects for sex (*F*_(1, 28)_ = 19.2, *P* = 0.0002) and genotype (*F*_(1, 28)_ = 11.2, *P* = 0.002) with a significant interaction (*F*_(1, 28)_ = 5.09, *P* = 0.03); *n* = 8 in all groups; Fig. 1f-h). Specifically, in male *Mapt^-^*^/-^ rats there was significantly enhanced LTD (31.8 ± 3.1%) relative to male WT rats (17.9 ± 2.7%; Šídák’s post hoc test: *P* = 0.0014). In contrast, there was no difference in LTD levels between female *Mapt^-^*^/-^ rats (15.3 ± 1.5%) and female WT rats (12.6 ± 2.4%; *P* = 0.8), and the magnitude of LTD also did not differ between sexes of WT rats (*P* = 0.9).

The enhanced LTD in male *Mapt^-^*^/-^ rats was not the result of changes to basal synaptic properties, as the input/output (I/O) curve relationships between fibre volley (FV) and stimulation intensity, and field excitatory postsynaptic potential (fEPSP) and FV, along with paired-pulse facilitation (**PPF**; a form of short-term plasticity), were all unchanged in both sexes of rats *Mapt^-^*^/-^ rats (Fig. 1i-k and Extended Data Fig. 3). Furthermore, hippocampal synaptic expression and phosphorylation levels of GluN2A, GluN2B, GluA1 and GluA2 were largely unaffected by tau deletion (Extended Data Fig. 4). We did however note elevated levels of GluN2A in female versus male WT rats, and a trend for the same difference between sexes of tau knockouts (Extended Data Fig. 4a). In summary, the deletion of tau in rats results in a sex-specific enhancement of LTD without impacting basal synaptic function or the expression of key synaptic receptors.

### The impact of tau deletion on LTD is species dependent

Our finding of increased LTD in male *Mapt^-^*^/-^ rats contrasts with previous findings of reduced LTD in male *Mapt^-^*^/-^ mice^5^. Since LTD is sensitive to many factors, such as recording conditions, and stress^27^, we determined whether our findings represent a *bona fide* species difference by interleaving experiments using *Mapt^-^*^/-^ mice and a new cohort of rats. Consistent with the first cohort (Fig. 1f-h), we again observed enhanced LTD in male, but not female, *Mapt^-^*^/-^ rats relative to WT littermates in interleaved experiments with *Mapt^-^*^/-^ mice (Fig. 2). However, in agreement with the previous study^5^, we found that male *Mapt^-^*^/-^ mice exhibited reduced LTD compared to male WT mice (14.3 ± 3.5% versus 26.4 ± 1.6% LTD, respectively; Student’s *t*-test: *t*_(10)_ = 3.2, *P* = 0.01; *n* = 6 in both groups; Fig. 2a, b). We also noted a sex-difference in mice since there were no differences in LTD between female WT and *Mapt^-^*^/-^ mice (Fig. 2e, f). Thus, tau deletion has opposing effects on LTD levels in rats and mice, and these effects are male-specific.

**Figure 2.**
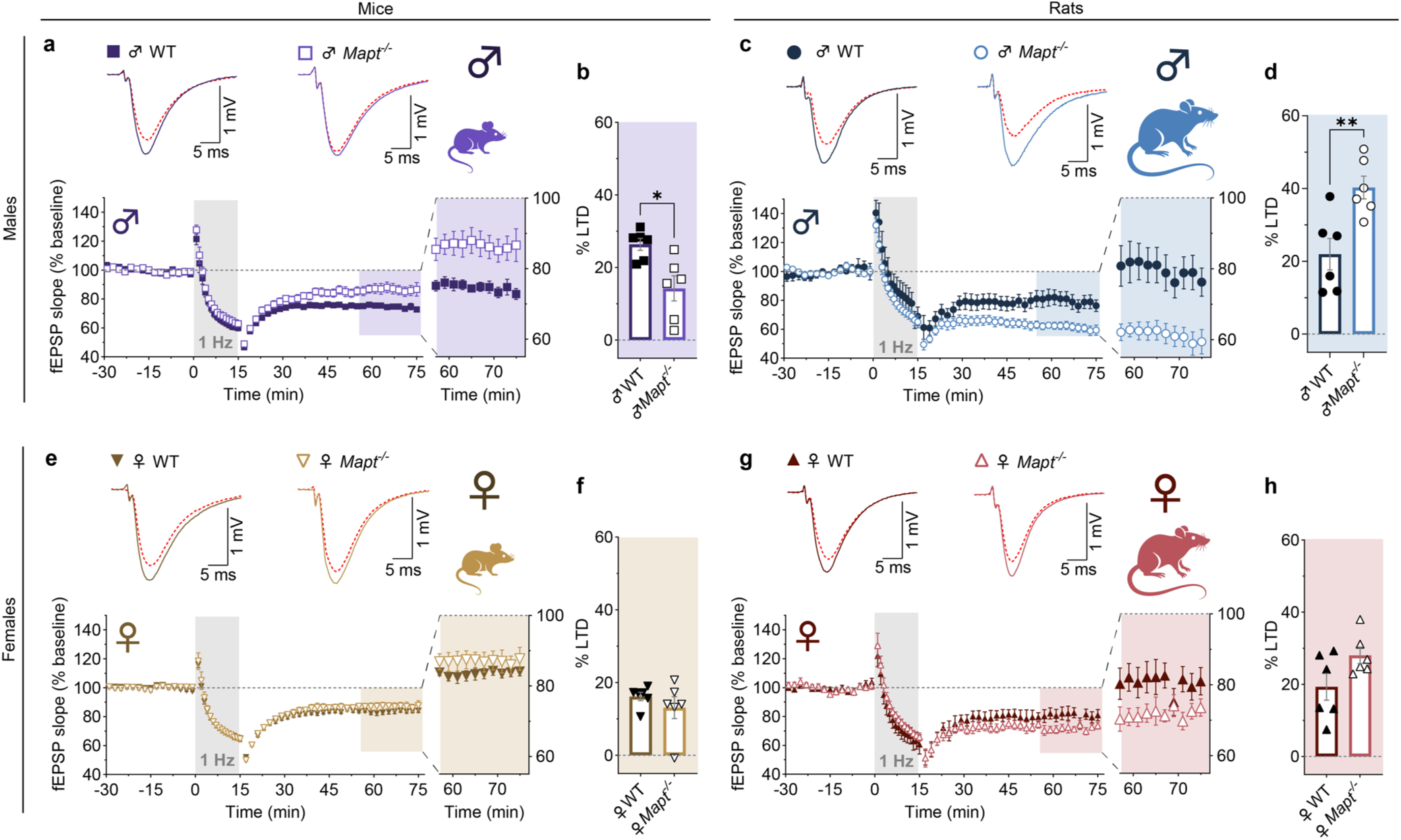
Opposite effect of *Mapt* deletion on LTD in male rats *versus* mice. **a**,**b**, LTD induced by LFS was reduced in male *Mapt^-/-^* mice (14.3 ± 3.5%) relative to male WT mice (26.4 ± 1.6%; *P* = 0.01). **c**,**d**, In interleaved recordings using a new cohort of rats, male *Mapt^-/-^* rats again exhibited significantly enhanced LTD (40.3 ± 3.1%) relative to male WT rats (22.0 ± 4.3%; *P* = 0.006). **e**,**f**, In contrast to males, LTD levels in female *Mapt^-/-^* mice (12.9 ± 4.7%) were similar to those observed in female WT mice (15.2 ± 1.6%; *P* = 0.4). **g**,**h**, Similarly, LTD levels in female *Mapt^-/-^* rats (28.1 ± 2.3%) did not differ significantly from those observed in female WT rats (19.3 ± 3.7%; *P* = 0.07). Expanded regions of the time plots show the final 20 min of recordings. Representative traces show superimposed fEPSPs from the baseline period (solid line) and the end of the recording session (dashed line). Levels of LTD were calculated as the percent change from baseline in the last 6 min of recordings. Comparisons were made using unpaired two-tailed Student’s *t*-tests (*n* = 6 in all groups). Detailed statistics are shown in Supplementary Table 2. \**P* < 0.05; \*\**P* < 0.01.

### Enhanced LTD in male *Mapt^-^*^/-^ rats is independent of NMDARs

We next sought to identify the receptor responsible for enhanced LTD in male *Mapt^-^*^/-^ rats. Since LTD induced by 1 Hz stimulation normally relies solely on the synaptic activation of NMDARs^17,28^, we tested the effects of the NMDAR glutamate site competitive antagonist D-AP5. As expected, inhibition of NMDARs eliminated LTD in WT male rats (0.1 ± 2.5%), as well as both WT (0.8 ± 2.6%) and *Mapt^-^*^/-^ (0.9 ± 3.1%) female rats. However, in the presence of D-AP5 there was a residual LTD in male *Mapt^-^*^/-^ rats (12.7 ± 1.9%) that significantly differed from male WT rats (Šídák’s post hoc test: *P* = 0.005; *n* = 8 in each group; Fig. 3). Therefore, it can be concluded the enhanced component of LTD in male rats is fully independent of NMDAR activation.

**Figure 3.**
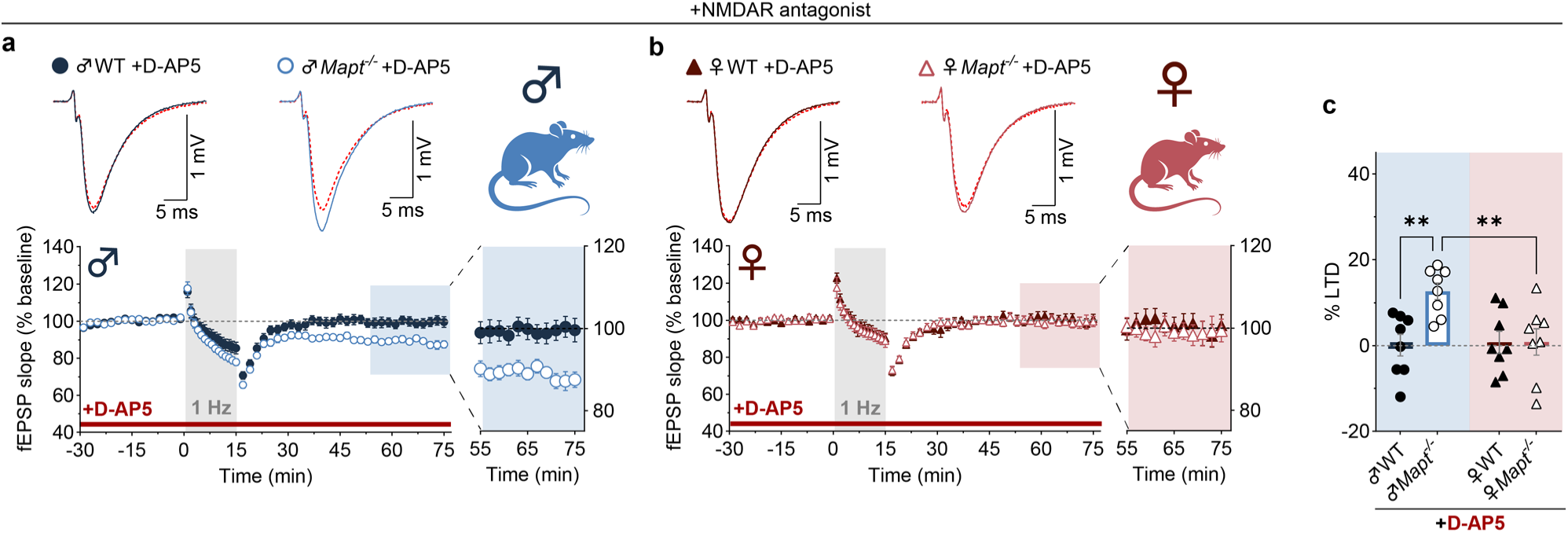
Residual NMDAR-independent LTD in male *Mapt^-/-^* rats. a-c, Effect of NMDAR blockade by D-AP5 (50 µM) on LFS-induced LTD in male and female WT and *Mapt*^-/-^ rats. LTD was fully blocked by D-AP5 in slices from male WT rats (0.1 ± 2.5%), whereas a residual NMDAR-independent LTD was apparent in slices from male *Mapt*^-/-^ rats (12.7 ± 1.9%; *P* = 0.005 relative to male WT) (**a**,**c**). In contrast, LTD was fully blocked by D-AP5 in slices from both WT (0.8 ± 2.6%) and *Mapt*^-/-^ (0.9 ± 3.1%) female rats (**b**,**c**). Expanded regions of the time plots show the final 20 min of recordings. Representative traces show superimposed fEPSPs from the baseline period (solid line) and the end of the recording session (dashed line). Levels of LTD were calculated as the percent change from baseline in the last 6 min of recordings. Comparisons were made using two-way ANOVA followed by Šídák’s post hoc test (*n* = 8 in all groups). Detailed statistics are shown in Supplementary Table 2. \*\**P* < 0.01.

### mGluR_I_s mediate the enhanced LTD in male *Mapt^-^*^/-^ rats

Under certain conditions or patterns of stimulation, mGluR_I_s can also mediate LTD at hippocampal CA1 synapses^16,29–31^. We therefore next investigated whether the residual NMDAR-independent LTD in male *Mapt^-^*^/-^ rats was dependent on mGluR_I_ activation. Since mGluR_I_-LTD induced at CA1 synapses can involve both mGluR_I_ subtypes mGlu1 and mGlu5^16^, we applied a combination of YM 298198 (**YM**) and MTEP to inhibit both of these receptors, respectively. In the presence of D-AP5, the addition of YM and MTEP abolished the residual LTD in male *Mapt^-^*^/-^ rats (Fig. 4a-c). We therefore next assessed the effects of mGluR_I_ antagonism in the absence of D-AP5. In control experiments, we again observed enhanced LTD in male *Mapt^-^*^/-^ (33.8 ± 2.9%) compared to WT (22.4 ± 2.6%) rats in a third cohort (Šídák’s post hoc test: *P* = 0.03; *n* = 9 in all groups; Fig. 4d,f). The addition of YM plus MTEP significantly reduced the enhanced LTD in *Mapt^-^*^/-^ rats (21.8 ± 3.7%; *P* = 0.02) but had no effect on LTD in WT rats (27.1 ± 1.3%; *P* = 0.7), relative to *Mapt^-^*^/-^ and WT controls, respectively (Fig. 4e,f). As such, the level of LTD in male *Mapt^-^*^/-^ rats in the presence of YM plus MTEP was similar to that observed in male WT rats (*P* = 0.6). Taken together, these experiments indicate that the enhanced component of LTD in male *Mapt^-^*^/-^ rats requires mGluR_I_ signaling.

**Figure 4.**
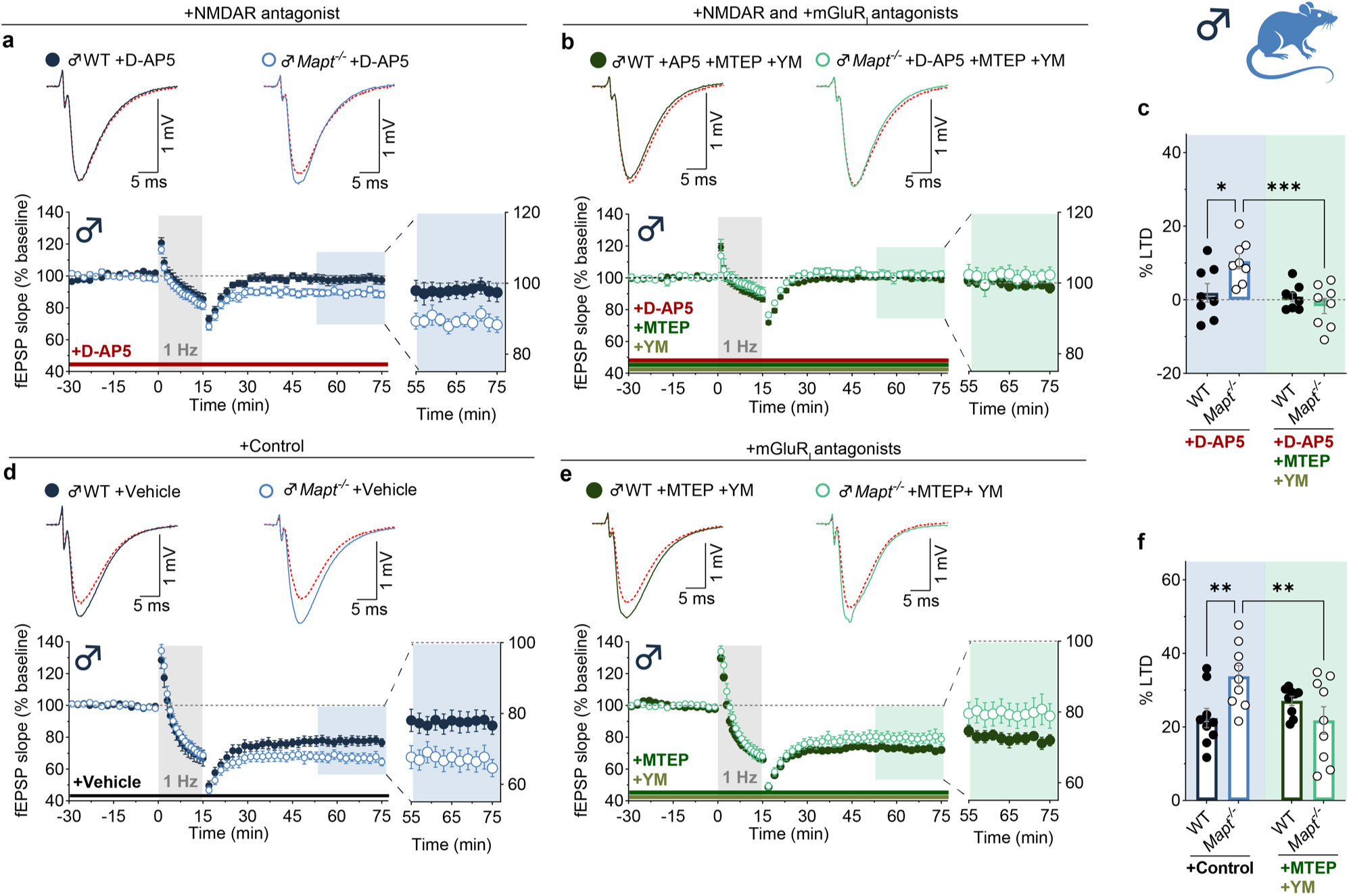
Enhanced LTD in male *Mapt^-/-^* rats is mGluR_I_-dependent. **a**-**c**, Effect of mGluR_I_ antagonism by YM (1 µM) and MTEP (1 µM) on the residual LTD induced in male *Mapt*^-/-^ rats in the presence of D-AP5 (50 µM). In control recordings (D-AP5 alone), LFS again induced a residual NMDAR-independent LTD in male *Mapt*^-/-^ rats (10.5 ± 2.0%) relative to the full block of LTD observed in male WT rats (1.9 ± 2.5%, *P* = 0.002) (**a**,**c**). The addition of YM+MTEP abolished the residual LTD in male *Mapt*^-/-^ rats (−1.7 ± 2.1%; *P* = 0.0008 relative to *Mapt*^-/-^ D-AP5 control) while having no additional effect on LTD levels in male WT rats (0.7 ± 1.2%; *P* = 0.9 relative to WT D-AP5 control) (**b**,**c**). **d**-**f**, Recordings (interleaved with those shown in **a**-**c**) showing the effect of YM+MTEP on LTD in male WT and *Mapt*^-/-^ rats in the absence of D-AP5. Enhanced LTD was again observed in male *Mapt*^-/-^ rats (33.8 ± 2.9%) relative to male WT rats (22.4 ± 2.6%, *P* = 0.03) in control recordings (**d**,**f**). mGluR_I_ antagonism had no effect on LTD in male WT rats (27.1 ± 1.3%; *P* = 0.7 relative to WT control) but reduced the level of LTD in male *Mapt*^-/-^ rats (21.8 ± 3.7%; *P* = 0.02 relative to *Mapt*^-/-^ control) such that it did not differ significantly from the LTD levels observed in male WT rats in the presence YM+MTEP (*P* = 0.6) (**e**,**f**). Expanded regions of the time plots show the final 20 min of recordings. Representative traces show superimposed fEPSPs from the baseline period (solid line) and the end of the recording session (dashed line). Levels of LTD were calculated as the percent change from baseline in the last 6 min of recordings. Comparisons were made using two-way ANOVA followed by Šídák’s post hoc test (*n* = 9 in all groups). Detailed statistics are shown in Supplementary Table 2. \**P* < 0.05; \*\**P* < 0.01.

### Enhanced LFS-LTD is not due to altered mGluR_I_ expression and is distinct from DHPG-LTD

The simplest explanation for an involvement of mGluR_I_s in LFS-induced LTD is an increase in synaptic mGluR_I_ levels. However, we found no significant differences in mGlu1 and mGlu5 dimer or monomer levels in either synaptically-enriched or in total hippocampal homogenates in either sex of *Mapt^-^*^/-^ rats relative to sex-matched WT rats (Extended Data Fig. 5a-l). We also assayed a form of mGluR_I_-dependent LTD induced by the mGluR_I_ agonist (*S*)-3,5-dihydroxyphenylglycine (**DHPG**)^32,33^, which acts on the total pool of surface (both synaptic and extrasynaptic) mGluR_I_s. DHPG-LTD was not significantly different between WT and *Mapt^-^*^/-^ rats of either sex (Extended Data Fig. 5m-o). Therefore, these results suggest that the enhanced LFS-LTD in male *Mapt^-^*^/-^ rats is not due to increased expression of mGluR_I_s or alterations in signaling mechanisms downstream of mGluR_I_s that underlie DHPG-LTD.

### Actin depolymerization enhances LTD in WT, but not *Mapt^-^*^/-^, male rats

The activation of mGluR_I_s by synaptically released glutamate is sensitive to receptor subsynaptic localization^34^, which in turn is influenced by their association with binding partners including cytoskeletal components such as microtubules and actin^35,36^. In line with a role of tau in stabilizing the actin cytoskeleton^37^, we found that levels of β/γ-actin were significantly increased in synaptically-enriched, but not total, hippocampal homogenates prepared from both male *Mapt^-^*^/-^ rats (131.9 ± 12.8% of male WT; *P* = 0.03) and female *Mapt^-^*^/-^ rats (135.9 ± 8.4% of female WT; *P* = 0.01; Šídák’s post hoc test; *n* = 8 in all groups; Fig. 5b,d). In contrast, α-tubulin levels were unchanged in both synaptically-enriched and total hippocampal fractions of both sexes (Fig. 5a,c). These findings are indicative of a possible role for tau in regulating synaptic actin content.

**Figure 5.**
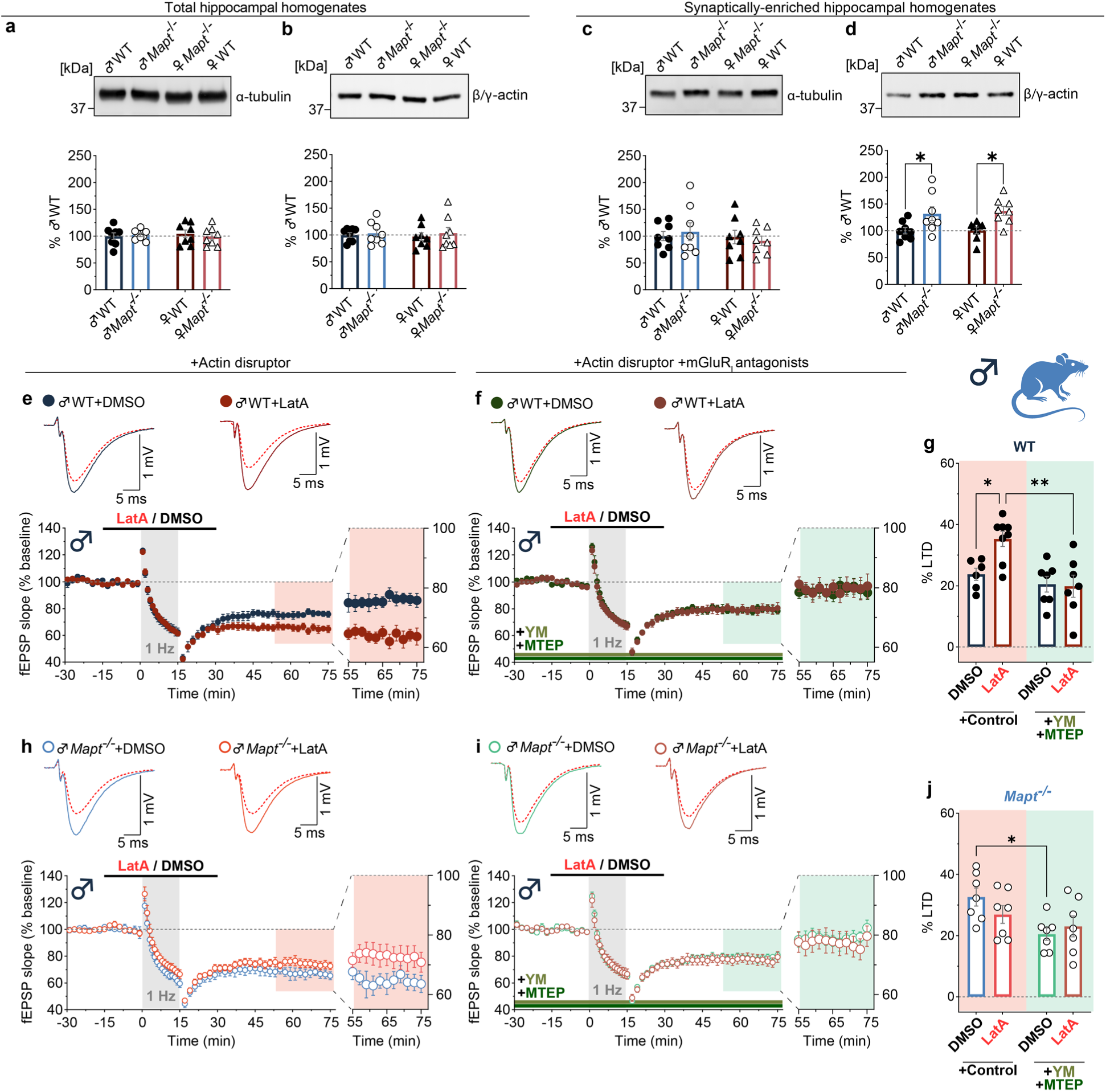
*Mapt* deletion increases synaptic β/γ-actin levels and occludes the mGluR_I_-dependent enhancement of LFS-LTD by latrunculin-A in male rats. **a-d**, Representative immunoblots (top) and quantification (bottom) of α-tubulin and β/γ-actin levels in total (**a**,**b**) and synaptically enriched (**c**,**d**) hippocampal homogenates from juvenile male and female WT and *Mapt^-/-^* rats. Both total and synaptic levels of α-tubulin (**a**,**c**), as well as total β/γ-actin levels (**b**), were unchanged in *Mapt^-/-^* rats of both sexes. However, synaptic levels of β/γ-actin (**d**) were increased in male *Mapt^-/-^*rats (131.9 ± 12.8%) relative to male WT rats (100.0 ± 5.9%; *P* = 0.03) and in female *Mapt^-/-^*rats (137.2 ± 8.5%) relative to female WT rats (100.9 ± 6.0%; *P* = 0.01, *n* = 8 in all groups). **e**-**j**, Interleaved recordings showing the effect of latrunculin A (LatA; 0.1 µM), an inhibitor of actin polymerization, on LFS-LTD in male WT and *Mapt^-/-^* rats, either in the absence (**e**,**h**; control) or presence (**f**,**i**) of the mGluR_I_ antagonists YM (1 µM) and MTEP (1 µM). In the control condition, treatment of male WT slices with LatA prior to, and during, delivery of LFS (as indicated by the bar in the figure) resulted in a significant enhancement of LTD (35.3 ± 2.5%, *n* = 8) relative to DMSO (0.001%) controls (23.8 ± 2.0%, *n* = 6; *P* = 0.03) (**e**,**g**). This LatA-mediated LTD enhancement was blocked in the presence of YM+MTEP (19.9 ± 3.6%, *n* = 7; *P* = 0.002 relative to LatA control), whereas mGluR_I_ antagonism had no effect on LTD in DMSO-treated slices from male WT rats (20.6 ± 2.6%, *n* = 7; *P* = 0.9 relative to DMSO control). In contrast to male WT slices, LatA treatment alone had no significant effect on LTD (26.9 ± 2.9%, *n* = 7) relative to DMSO controls (32.6 ± 2.9, *n* = 7; *P* = 0.5) in slices from male *Mapt^-/-^* rats (**h**,**j**). Similarly, LTD levels in LatA-treated male *Mapt^-/-^* slices (20.5 ± 2.0%, *n* = 7) did not differ from those in DMSO-treated slices (23.0 ± 3.4%, *n* = 7; *P* = 0.8) under conditions of mGluR_I_ antagonism (**i**,**j**). However, the enhanced LTD present in male *Mapt^-/-^* rats was significantly reduced by YM+MTEP in DMSO-treated slices (*P* = 0.03). Expanded regions of the time plots show the final 20 min of recordings. Representative traces show superimposed fEPSPs from the baseline period (solid line) and the end of the recording session (dashed line). Levels of LTD were calculated as the percent change from baseline in the last 6 min of recordings. Comparisons were made using two-way ANOVA followed by Šídák’s post hoc test. Detailed statistics are shown in Supplementary Tables 2 and 3. \**P* < 0.05; \*\**P* < 0.01.

To investigate more directly whether actin is functionally involved in the enhanced mGluR_I_-dependent LTD identified in male *Mapt^-^*^/-^ rats, we treated hippocampal slices with Latrunculin A (**LatA**) prior to, and during, LFS. Latrunculin compounds sequester globular actin to prevent polymerization, resulting in the depolymerization of actin filaments^38^ and the modulation of synaptic plasticity and G/F actin ratio^39–41^. In line with this, we found that LatA enhanced LTD in male WT rats relative to DMSO controls (35.3 ± 2.5% versus 23.8 ± 2.0%; Šídák’s post hoc test: *P* = 0.03; *n* = 8 and 6, respectively; Fig. 5e,g). Moreover, this LatA-mediated enhancement of LTD was prevented when mGluR_I_ activation was blocked by the presence of YM plus MTEP in the perfusate (19.9 ± 3.6% LTD; *P* = 0.002 relative to LatA alone; n = 7; Fig. 5f,g). In contrast, LTD levels in male *Mapt^-^*^/-^ rats (32.6 ± 2.9% in DMSO controls, *n* = 7) were not enhanced by LatA treatment (26.9 ± 2.9%; Šídák’s post hoc test: *P* = 0.5; *n* = 7; Fig. 5h,j), but they were again reduced by mGluR_I_ antagonism (20.5 ± 2.0%; *P* = 0.03; *n* = 7; Fig. 5i,j), suggesting that the absence of tau mimics and occludes the enhancement of LTD by LatA. Furthermore, and in contrast to males, LatA had no effect on LTD in either WT or *Mapt^-^*^/-^ female rats (Extended Data Fig. 6). Together, these findings suggest that tau deletion alters the actin cytoskeleton, and this occurs in a manner that influences mGluR_I_ activation during LFS-LTD in male but not female rats.

## DISCUSSION

In the present study we have identified species-and sex-dependent effects of tau deletion on LTD in the CA1 region of the hippocampus. Specifically, using a new *Mapt^-^*^/-^ rat model, we observed that juvenile male rats lacking tau have enhanced LFS-induced LTD relative to sex-matched WT littermates. In contrast, male *Mapt^-^*^/-^ mice exhibited reduced LTD, consistent with previous reports^5^. Interestingly, LTD in females was unaffected by tau elimination in both rats and mice. Consistent with previous studies^17,28,42^, the form of LTD that was induced by LFS in WT rodents was fully dependent on the synaptic activation of NMDARs. However, the additional LTD observed in male *Mapt^-^*^/-^ rats required the synaptic activation of mGluR_I_s but not NMDARs. The enhanced LTD was not associated with any alterations in the total surface expression of mGluR_I_s since their pharmacological activation, using DHPG, elicited similar levels of LTD in WT and *Mapt^-^*^/-^ rats. This suggests that a sex-dependent sub-synaptic redistribution of mGluR_I_s from perisynaptic/extrasynaptic to synaptic sites may underlie the effect. Indeed, tau deletion increased synaptic β/γ-actin levels in juvenile *Mapt^-^*^/-^ rats, pointing to a role of the actin cytoskeleton. Furthermore, tau deletion mimicked the effect of LatA on LTD in male WT rats^39^, whereas the ability of LatA to enhance LTD was occluded in male *Mapt^-^*^/-^ rats. Both manipulations enhanced LTD in a manner that was fully dependent on the synaptic activation of mGluR_I_s. Together, our data reveal an unexpected sex-specific role of tau in regulating the involvement of mGluR_I_s during LTD in rats, via an interaction with the synaptic actin cytoskeleton.

### Generation of a *Mapt* knockout rat model

Our interest in making a tau knockout rat was to investigate the physiological synaptic function of tau in a species that has a greater cognitive repertoire than mice^43,44^. Tau is the principal protein that becomes dysregulated in tauopathies, including AD. It is noteworthy, that rats display more robust signs of tau pathology in AD models compared to mice^45,46^ and have different developmental expression profiles for tau isoforms. For example, in the CNS, 1N3R and 2N3R tau isoforms persist throughout the lifespan at low levels in rats but are expressed only during early development or are undetectable in mice^47–51^. Although the present study does not address the function of tau isoforms, the rat tau knockout is a useful tool to identify the isoform-specific functions of endogenous and pathological tau through re-expression studies. Recently, another tau knockout rat (Cy23) was generated and found to extend survival when crossed with the Tg12099 (tau *0N4R MAPT*P301S*) model of frontotemporal dementia^20^. Consistent with the Cy23 rat line, we found no gross phenotypic differences in our *Mapt^-^*^/-^ rats. However, we studied LTD, a model for the early stages of synaptic loss^67^, and there were clear sex-dependent changes within our tau knockouts. This has implications for both the physiological function of synaptic tau, and potential consequences for tau-lowering therapies.

### Species-dependent effects of tau deletion on LTD in male rodents

We were surprised to observe that tau deletion enhances LTD in male rats, since previous work reported a substantial reduction in LTD in male mice^5^. In interleaved experiments using male rats and mice we confirmed that this is a *bona fide* species difference rather than some experimental difference, such as housing or recording conditions. We also found that, as in female rats, LTD in female mice was unaffected by tau deletion. The reason for this male-specific rodent species difference remains unknown and could be attributable to several factors. One plausible explanation is the lack of compensatory changes in other MAPs in our *Mapt^-^*^/-^ rats, which contrasts with juvenile *Mapt^-^*^/-^ mice^54^. A second possibility is different functions of tau that may arise^55^ due to tau isoform or expression differences between the two species^47–51^. A third possibility is extrinsic factors, such as differences in how rats and mice respond to stress^46^, for which tau has been shown to play a role in mice^56^. A fourth is intrinsic factors, such as the expression levels of synaptic molecules^57^, or the underlying mechanisms of LTD induction or expression that may differ between these rodent species^58^.

### mGluR_I_s mediate enhanced LTD in male *Mapt^-^*^/-^ rats

To assess the contribution of mGluR_I_s to the enhanced LTD in male *Mapt^-^*^/-^ rats, we used a combination of mGlu1 (YM) and mGlu5 (MTEP) antagonists, since the effects of mGluR_I_ activation at CA1 synapses may be mediated by an mGlu1/mGlu5 heterodimer that requires both subunits to be inhibited^16,29,30,59^. Typically, to induce mGluR_I_-LTD at these synapses, higher stimulation frequencies (e.g., 5 Hz)^30^ or paired-pulse LFS^60,61^ are required, suggesting that the absence of tau in male rats may result in a shift in the activation frequency for synaptically-induced mGluR_I_-LTD. Unlike LFS-induced LTD, DHPG-LTD was unaffected by tau deletion in male rats. This suggests that the total surface expression of mGluR_I_s is not altered in *Mapt^-^*^/-^ rats. Therefore, it is likely that deletion of tau results in increased synaptic mGluR_I_ function due to changes in the sub-synaptic localization of the receptors^34^.

### Involvement of the synaptic actin cytoskeleton in the mGluR_I_-dependent enhancement of LTD

To address how the absence of tau might impact synaptic mGluR_I_ activation, we investigated synaptic levels of two key tau binding partners – actin and tubulin. Previous studies have shown that tau regulates the synaptic cytoskeleton through its interaction with actins^24,62^, and the crosslinking of actin with microtubule networks^37^. In the present work, we found increases in β/γ-actin, but not α-tubulin, in synaptically-enriched hippocampal fractions from *Mapt^-^*^/-^ rats. This suggests that alterations in the synaptic actin cytoskeleton is a factor underlying the effect of tau deletion on LTD in male rats. Indeed, actin co-localizes with mGluRs, controls mGluR trafficking, and regulates LTD^34,36,40^. Direct evidence for a role of the synaptic actin skeleton is provided from the findings that LatA enhanced LTD in male, but not female, WT rats, while having no additional effect in male *Mapt^-^*^/-^ rats. Furthermore, the enhancements of LTD resulting from both LatA treatment and tau deletion were dependent on activation of mGluR_I_s. We conclude, therefore, that enhanced LTD in male *Mapt^-^*^/-^ rats arises from an alteration in the actin-dependent localization of synaptic mGluR_I_s. The sex-specific effect of tau deletion on LFS-LTD may arise from the influence of sex hormones. Indeed, synaptic plasticity in the hippocampus of rodents^63^, including prepubescent rats^64^, has been shown to be influenced by estrogen receptors, which are known to impact plasticity via the actin cytoskeleton^65^.

### Implications for the understanding of AD and other tauopathies

Although the purpose of the present investigation is to understand the physiological function of synaptic tau in juvenile animals, the observations have broader implications for neurodegenerative diseases involving tau. In particular, acutely delivered β-amyloid (**Aβ**) and tau species can rapidly impact synaptic function, suggesting that these molecules may trigger neurodegeneration in advance of the formation of plaques and tangles^66^. Potentially, neurodegeneration may be initially triggered by the local build up of toxic Aβ species leading to hyperphosphorylation of tau, via a mechanism triggered by mGluR_I_s and involving glycogen synthase kinase 3 (**GSK-3**)^67,68^, and activation of the complement cascade^69^.

Pertinent to the present study is the finding that Aβ binds to mGlu5 receptors in male but not female humans and mice, via a cellular prion protein (PrP^c^)-dependent mechanism^70,71^. Critically, this pathological cascade can be mitigated, as pharmacological inhibition of mGlu5 can reverse cognitive deficits in male but not female adult AD rodents, by alleviating aberrant GSK-3 activation and autophagy impairments^72^. Therefore, the increase in synaptic weakening triggered by mGluR_I_s, as defined in the present study, could underlie this cognitive dysfunction in males. In females, however, a different mechanism is likely to underlie the neuropathology. Indeed, cognitively impaired female, but not male, patients exhibit lower mGlu5 levels associated with greater tau deposition^73^.

### Conclusions

Most studies on tau have focussed on its seeding, propagation, and aggregation. However, increasing evidence has shown that dysregulation of tau precedes these critical factors and is associated with early cognitive dysfunction. Therefore, understanding these early changes is of direct relevance for both understanding the aetiology of tauopathies and to the design of effective therapeutics. To achieve this effectively, it is first necessary to define the physiological function of tau at synapses. We have shown how tau regulates synaptic plasticity, and since this occurs early in development it can be considered a physiological function of tau. The enhancement of mGluR_I_ function in male *Mapt^-^*^/-^ rats provides a rational explanation for the effectiveness of mGluR_I_ antagonists in experimental models of AD in males^72^. These findings are also relevant to the understanding of how tau loss-of-function contributes to tauopathies^74^, and for understanding the effects of tau-lowering therapies in humans^75,76^. The availability of the *Mapt^-^*^/-^ rat should now enable the mechanistic basis for female cognitive decline to be established and an explanation for the increased susceptibility of females to be obtained.

## METHODS

### Creation of *Mapt^-/-^* rat line

The constitutive *Mapt^-/-^* rat line (SD-*Mapt*^em1Tcp^) was generated by delivering CRISPR-Cas9 reagents directly to Charles River Laboratories caesarean-derived Sprague Dawley (Crl:CD(SD)) rat zygotes, following established protocols^22^. Sprague-Dawley (#001) rats were purchased from Charles River Laboratories (Montreal, QC) and maintained in the Canadian Council on Animal Care (CCAC)-accredited animal facility at The Hospital for Sick Children (Toronto) in individually ventilated cages (Allentown) on a 12-h light/dark cycle (7 am on/7 pm off). Preparation of embryo donors and recipients, as well as embryo transfer surgeries were based on previously described procedures^77^. Four guide RNAs were synthesized using EnGen sgRNA Synthesis Kit (New England BioLabs, E3322) and were combined with *Streptococcus pyogenes* and 656 ng/µL of Cas9 protein (Integrated DNA Technologies, 1074182). Electroporation was performed as previously described^78^, but with modified electroporation parameters. Zygotes were briefly washed through Acidic Tyrode’s solution (Sigma T1788) and then Opti-MEM media (Thermo Fisher Scientific 31985062). The zygotes were then placed into 20 μL of Cas9 RNP mix and the whole volume was transferred to 1 mm-gap cuvette. Twelve square pulses at 30V with 3 ms pulse duration and 100 ms interval were applied using BioRad Gene Pulser XCell electroporator. The zygotes were retrieved from the cuvette with a Rainin pipettor and 20 µL tips followed by at least two rinses of 70 μL of Life Global media (LGGG-050, Cooper Surgical). Electroporated zygotes were implanted into pseudopregnant SD females. Founder progenies were screened for the desired PCR product (823 – 985 base pairs), and deletion of the targeted *Mapt* gene region was confirmed using Sanger sequencing (Extended Data Fig. 1a). The deletion was predicted to introduce a premature stop codon, preventing protein expression of all tau isoforms. Tau protein knockout was confirmed in whole brain lysates using an *N*-terminal specific monoclonal tau antibody (Extended Data Fig. 1b).

### Animals

Juvenile (postnatal day (P) 14-to-19) wild-type (**WT**) and *Mapt^-/-^*rats (**SD-*Mapt*^em1Tcp^**) and mice (B6.129X1-*Mapt*^tm1Hnd^/J)^79^ of both sexes were used in this study. Heterozygous breeding pairs (*Mapt^+/-^* × *Mapt^+/-^*) were maintained at University Health Network (UHN; Toronto) facilities and replaced every 6-months. All experiments were conducted in compliance with UHN animal use protocols, with oversight from The Centre for Phenogenomics (TCP; Toronto). To maintain the genetic line, *Mapt^-/-^* rats and mice were backcrossed every 10 generations. Animals were housed under standard conditions on a 12-h light/dark cycle with *ad libitum* access to food and water. All experiments were conducted blind to genotype and utilized sex-matched WT and *Mapt^-/-^* rat or mouse littermate pairs for between-genotype comparisons. Unblinding occurred after completing experiments and data analysis.

### Genotyping

Tail or ear notch samples were collected from P8-to-12 old rats and mice. Genotyping was performed using an established NaOH extraction protocol^80^. Briefly, 75 µL of 25 mM NaOH/0.2 mM EDTA was added to each sample and heated at 95°C for 15 min. Samples were then centrifuged at 4,000 rpm for 3 min, and 75 µL of 40 mM Tris HCl at pH = 5.5 was added to neutralize the reaction.

For polymerase chain reaction (PCR), 1 µL of lysate was mixed with 5 µL of 2 × HS-red Taq mix, 3 µL of ultrapure water, and 1 µL of primers (final concentration: 4 µM). PCR was performed on a T-100 Thermal Cycler for 35 cycles (in min × °C): 5 × 95, 0.25 × 95, 0.25 × 65, and 0.5 × 72. Samples were then cooled and maintained on a cycle (in min × °C): 1 × 72, and ∞ × 12. PCR products were loaded onto a 2 % agarose gel in 1 × tris-acetate-EDTA buffer and subjected to electrophoresis at 180 V for 30 min in a Sub-Cell GT System (Bio-Rad). Gels were imaged using a GelDoc XR+ imaging system.

### Hippocampal slice preparation

Animals were euthanized by decapitation under deep isoflurane anesthesia. Brains were rapidly extracted and placed in chilled oxygenated (95% O_2_/5% CO_2_) artificial cerebrospinal fluid (ACSF) for 3-to-5 min. The dissection solution was the same composition as the recording ACSF, consisting of (in mM): 124 NaCl, 10 D-Glucose, 24 NaHCO_3_, 3 KCl, 1.25 NaH_2_PO_4_, 1 MgSO_4_, and 2 CaCl_2_. The hippocampus was extracted from both hemispheres, and transverse slices of dorsal hippocampus (400 µm thick) were prepared using a vibratome (model VT1200, Lecia Biosystems). The CA3 region was not removed from slices. Slices were recovered at room temperature for 1-2 hours before commencing experiments. No experiments were started beyond 6-hours following collection of slices.

### Extracellular hippocampal slice electrophysiology

Hippocampal slices were placed in a submerged recording chamber and continuously perfused with ACSF at a rate of 2.5 mL/min and maintained at 29°C. Fibre volleys (FVs) and field excitatory postsynaptic potentials (fEPSPs) were evoked by stimulating the Schaffer collateral-commissural pathway (SCCP) with a bipolar matrix stimulation electrode (model: MX21XEP(RM1), FHC). Responses were measured using a borosilicate glass recording electrode (1.3 – 4.0 MΩ) placed in the CA1 stratum radiatum approximately 300 µm away from the site of stimulation. Biphasic constant current stimulus pulses (0.1 ms duration primary pulse/0.2 ms duration half-intensity secondary pulse) were delivered using a stimulus isolating unit (STG 4002, Multichannel Systems) at a test frequency of 0.033 Hz. Baseline stimulus intensity was set at 3 × the minimum current required to evoke a visually detectable (∼0.1 mV) fEPSP (range: 12-36 µA). Signals were amplified and low-pass filtered at 3 kHz using a Model 3000 amplifier (A-M systems Inc.), digitized at 40 kHz using a USB-6341 digitizer (National Instruments), and acquired and analysed using WinLTP software^81^. Four consecutive stimulus sweeps were averaged for analysis, with FVs quantified by their peak amplitudes and fEPSPs quantified by their initial slopes. Input/output (I/O) curves were generated by evoking responses over five predetermined stimulus intensities (in µA): 0, 25, 50, 75, and 100. The I/O relationships between the stimulus intensity and FV amplitude, and the FV amplitude and fEPSP slope, were analyzed using linear regression, with the slope values of the linear fits used for statistical analysis. Paired-pulse facilitation (PPF) was assessed by delivering paired stimuli at 50, 100, 200, and 500 ms inter-pulse intervals. The paired-pulse ratio (PPR) was calculated by dividing the slope of the second fEPSP by the slope of the first fEPSP. Long-term depression (LTD) was induced after 30 min of stable baseline recording by delivering low-frequency stimulation (LFS) consisting of 900 pulses delivered at 1 Hz. DHPG-LTD was induced by applying 50 µM DHPG ((S)-3,5-Dihydroxyphenylglycine, Hello Bio, HB0045) for 10 min. The level of LTD in each experiment was quantified as the percent change in fEPSP slope from baseline in the final 6 min of recording (60 min post-induction).

### Pharmacological reagents

Compounds were prepared as stock solutions in ultrapure water or DMSO and diluted to final concentration in ACSF immediately prior to application via the perfusion system in hippocampal slice recordings. Final concentrations were as follows (in μM): 50 D-AP5 (D-2-amino-5-phosphonopentanoic acid, Hello Bio, HB0225), 1 YM 298198 hydrochloride (6-Amino-*N*-cyclohexyl-*N*,3-dimethylthiazolo[*3,2-a*] benzimidazole-2-carboxamide, Hello Bio, HB0664), 1 MTEP hydrochloride (3-((*2*-Methyl-*4*-thiazolyl)ethynyl)pyridine, Hello Bio, HB0431), and 0.1 Latrunculin A (Hello Bio, HB0375).

### Synaptic enrichment

Whole hippocampi from both hemispheres were extracted and snap frozen in liquid nitrogen. Synaptic enrichment was performed as outlined in^82^. Briefly, hippocampi were homogenized with 30 strokes in a Dounce homogenizer in 2 mL of high sucrose buffer (containing (in mM: 320 sucrose, 10 Tris-HCl (pH = 7.4), 1 EDTA, 1 EGTA) supplemented to a final 1 × concentration with protease and phosphatase inhibitor cocktail. 180 µL of homogenate was collected as the total hippocampus homogenate fraction and diluted with 20 µL of 10 × RIPA lysis buffer (Millipore, Cat #: 20-188). The remaining 1.82 mL of homogenate underwent centrifugation at 4°C, starting with 800 × g for 10 min. The supernatant was then pelleted at 9,200 × g for 15 min, washed with high sucrose buffer and centrifuged again under the same conditions. The final pellet underwent hypotonic lysis on ice in 2 mL of low sucrose buffer (in mM: 35.6 sucrose, 10 Tris-HCl (pH = 7.4), 1 EDTA, and 1 EGTA) with a final 1 × concentration of protease and phosphatase inhibitor cocktail. After centrifugation at 25,000 × g for 20 min at 4°C, the pellet was resuspended in 1 × RIPA lysis buffer. Protein concentrations were measured using the Pierce BCA Protein Assay Kit. Samples were reduced with β-Mercaptoethanol to a final concentration of 2.7% and stored at -80°C until use. Aliquots underwent no more than 3 × freeze/thaw cycles before being discarded.

### Immunoblotting

20 µg of protein was loaded onto 7.5% or 10% TGX Stain-Free FastCast polyacrylamide gels, with Precision Plus Protein All Blue Prestained Protein ladder added to one lane. Gels were placed in running buffer (1 × tris-glycine-SDS buffer and electrophoresed at 180 V until the desired separation was achieved. Total protein levels were tagged and imaged using stain-free technology (Bio-Rad) on a ChemiDoc MP imaging system.

Proteins were transferred onto Immun-Blot 0.45 µm low fluorescence PVDF membranes in transfer buffer (containing: 20% Trans-Blot Turbo 5 X Transfer Buffer, 20% ethanol, and 60% ultrapure water) across two cycles of Turbo-TGX transfer (25 V, 2.5 A, 3 min) on the Trans-Blot Turbo Transfer System. Membranes were rinsed in 1 × tris-buffered saline containing 0.1% Tween 20 (1 × TBST), and total protein levels were imaged. Blocking was performed using Everyblot blocking buffer (Bio-Rad, #120-100-20) for 30 min. Membranes were incubated overnight (16 h) at 4°C with primary antibodies (Supplementary Methods Table 1) on a gentle rocker. The following day, primary antibodies were removed, and the membranes were washed 6 times with 1 × TBST before and after incubation with secondary antibodies (Supplementary Methods Table 1) for 1 h at room temperature. Proteins levels were normalized to total stain-free protein levels and phosphorylated protein epitopes were normalized to total protein levels and expressed as a % relative to male WT rats.

### Data analysis and statistics

All reported *n*-values represent biological replicates (one slice/sample per animal) for all electrophysiology and biochemistry experiments. Data were graphed and underwent statistical analysis using GraphPad Prism Software (version 10.1.2) and are presented as the mean ± the standard error of the mean (s.e.m.). The statistical tests used are detailed in the results and/or figure legends, and the study data summary can be found in Supplementary Tables 1-3.

## ACKNOWLEDGEMENTS

The authors would like to thank the staff of the Model Production Core at The Centre for Phenogenomics (TCP) for excellent technical assistance during rat model production. We are grateful to Dr. Michael Salter and Janice Hicks for their initial support in maintaining the *Mapt^-^*^/-^ rat line. Portions of the study were published in the PhD thesis of L.T.R. (University of Toronto, 2024) and presented in conference abstracts.

The research was funded by a CIHR (Canadian Institutes of Health Research) Foundation Grant #154276 to G.L.C. and a Project Grant (PJT-173497) to P.F. We thank the Dani Reiss Family Foundation (Neurodegeneration and Aging Research Program) as well as Butch Mandel for their respective support. Generation of tau (SD-*Mapt*^em1Tcp^) knockout rat line was funded in part by Genome Canada and Ontario Genomics (OGI-137). G.L.C. is the holder of the Krembil Family Chair in Alzheimer’s Research. We also thank the Peterborough K.M. Hunter Charitable Foundation for providing funding support to L.T.R during this project.

## AUTHOR CONTRIBUTIONS

L.T.R., G.L.C., J.G., P.T., and E.W.S. wrote the manuscript and designed the experiments. L.T.R., with assistance from P.T., E.W.S, and L.A.J., performed and analyzed electrophysiology recordings. Immunoblotting experiments were performed and analyzed by B.P., M.W., L.T.R. and M.T. with support from J.S.T. and S.C. The *Mapt^-^*^/-^ rat line was generated by L.N. and M.G. Genotyping was performed by L.Z., A.K. and F.G. Project guidance was provided by G.S., Z.J., P.F., and C.A.B. All authors discussed the findings and edited the manuscript.

---

**Extended Data Figure 1.**
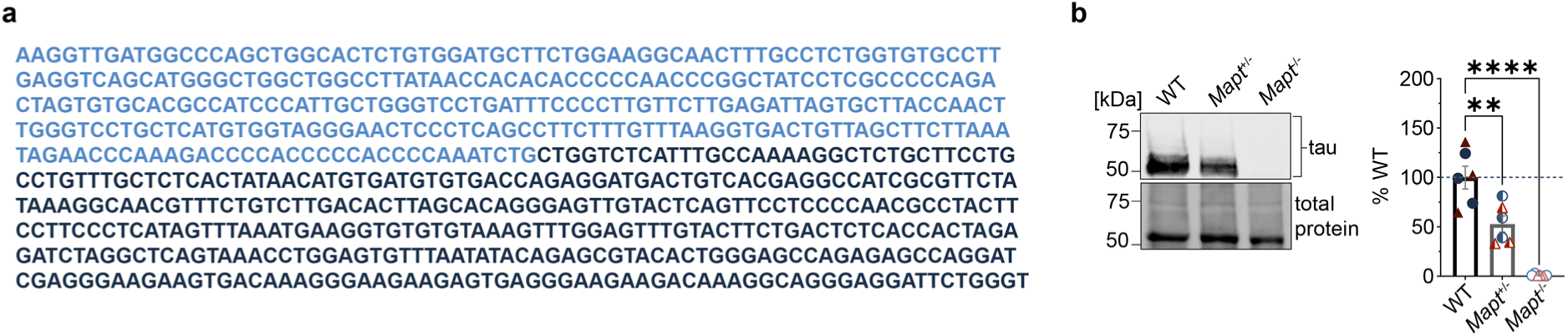
Molecular validation of SD-*Mapt*^em1Tcp^ rats. **a**, Results of Sanger sequencing of the *Mapt* gene in *Mapt^-/-^* rats showing the sequences upstream (light blue) and downstream (dark blue) of the deleted region. **b**, Representative immunoblot (left) and quantification (right) showing levels of tau expression in whole brain homogenates from juvenile male (circles) and female (triangles) *Mapt^+/+^* (WT), *Mapt^+/-^*, and *Mapt^-/-^* rats (*n* = 3 males and 3 females per group) using an *N*-terminal tau monoclonal antibody. Tau protein levels are normalized to total (stain-free) protein levels and expressed as % of WT (mean ± s.e.m.). \*\**P* < 0.01, \*\*\*\**P* < 0.0001.

**Extended Data Figure 2.**
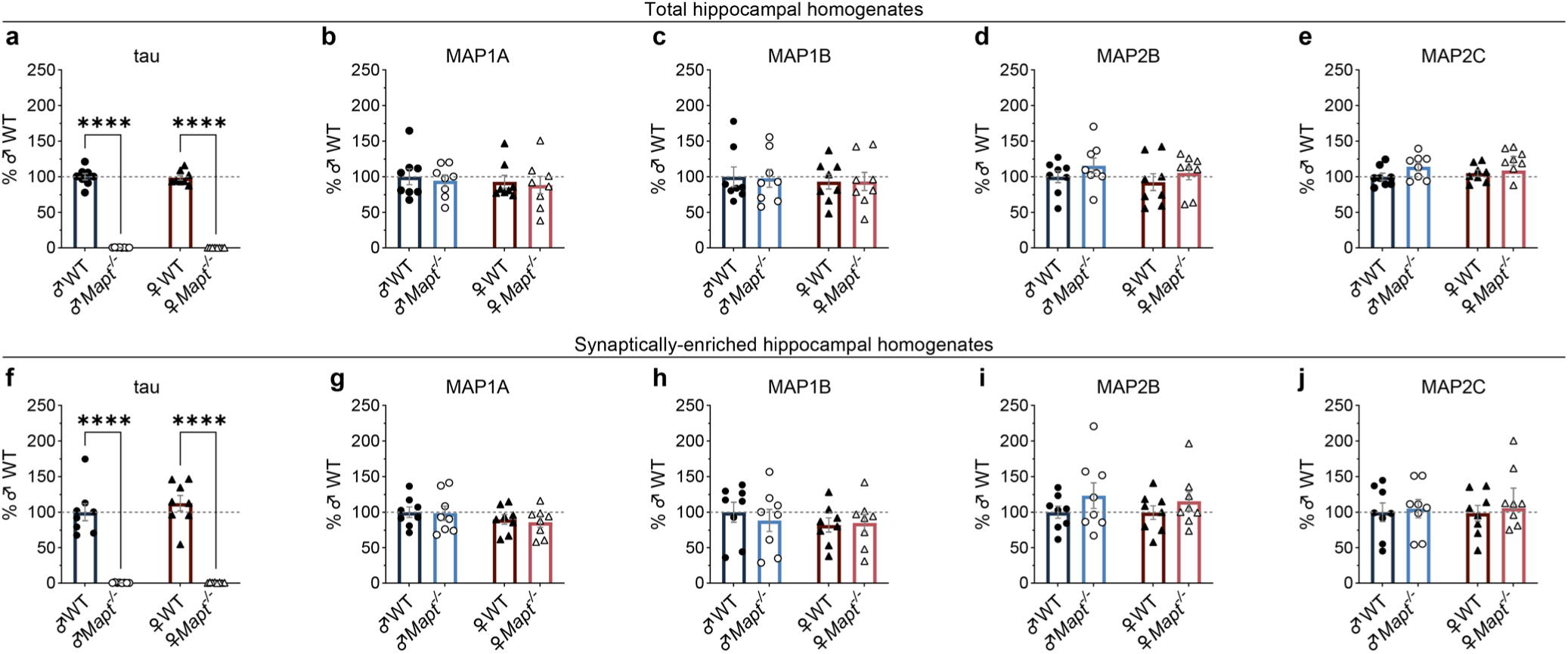
Total and synaptic hippocampal MAP expression levels in male and female WT and *Mapt^-/-^* rats. **a**-**m**, Quantification of tau, MAP1A, MAP1B, MAP2B, and MAP2C levels in total (**a**-**e**) and synaptically enriched (**f**-**j**) hippocampal homogenates from both sexes of WT and *Mapt^-^*^/-^ rats (*n* = 8 per group; representative blot images are shown in Fig. 1b,c). In total homogenates, male (0.5 ± 0.1%) and female (0.3 ± 0.03%) *Mapt^-^*^/-^ rats lacked tau protein relative to male (100.0 ± 4.4%; *P* < 0.0001) and female (98.6 ± 3.4%; *P* < 0.0001) WT rats, respectively (**a**). Tau was detectable in synaptically enriched hippocampal fractions from both male (100.0 ± 12.1 %) and female (112.7 ± 10.9%) WT rats but not male (0.6 ± 0.2%, *P* < 0.0001) and female (0.8 ± 0.2%, *P* < 0.0001) *Mapt^-^*^/-^ rats (**f**). Levels of all other MAPs in total homogenates (**b**-**e**) and synaptically enriched fractions (**g**-**j**) were not significantly altered in *Mapt^-^*^/-^ rats of either sex. Comparisons were made using two-way ANOVA followed by Šídák’s post hoc test. Detailed statistics are shown in Supplementary Table 3. \*\*\*\**P* < 0.0001.

**Extended Data Figure 3.**
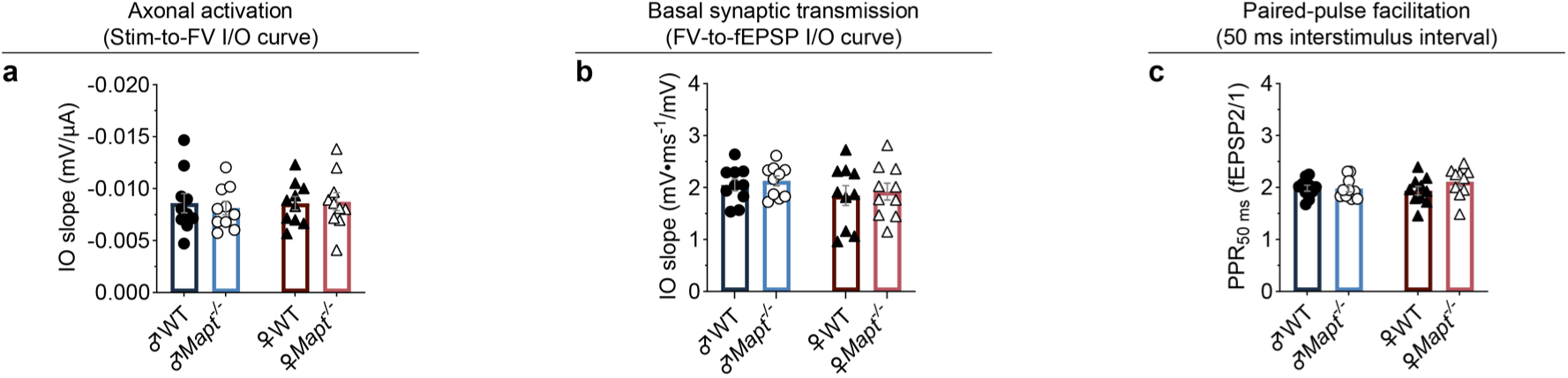
Quantification of I/O curve relationships and PPF in male and female WT and *Mapt^-/-^* rats. **a**,**b**, Linear regression slopes for the FV vs. stimulus intensity (**a**) and fEPSP vs. FV (**b**) I/O relationships. **c**, PPF at the 50 ms interstimulus interval, measured by calculating the paired-pulse ratio (PPR; fEPSP2/fEPSP1). There were no genotype or sex differences in any of these measures (*n* = 10 in all groups). Representative traces, I/O and PPF plots appear in Fig. 1i-k. Comparisons by two-way ANOVA. Detailed statistics are shown in Supplementary Table 2.

**Extended Data Figure 4.**
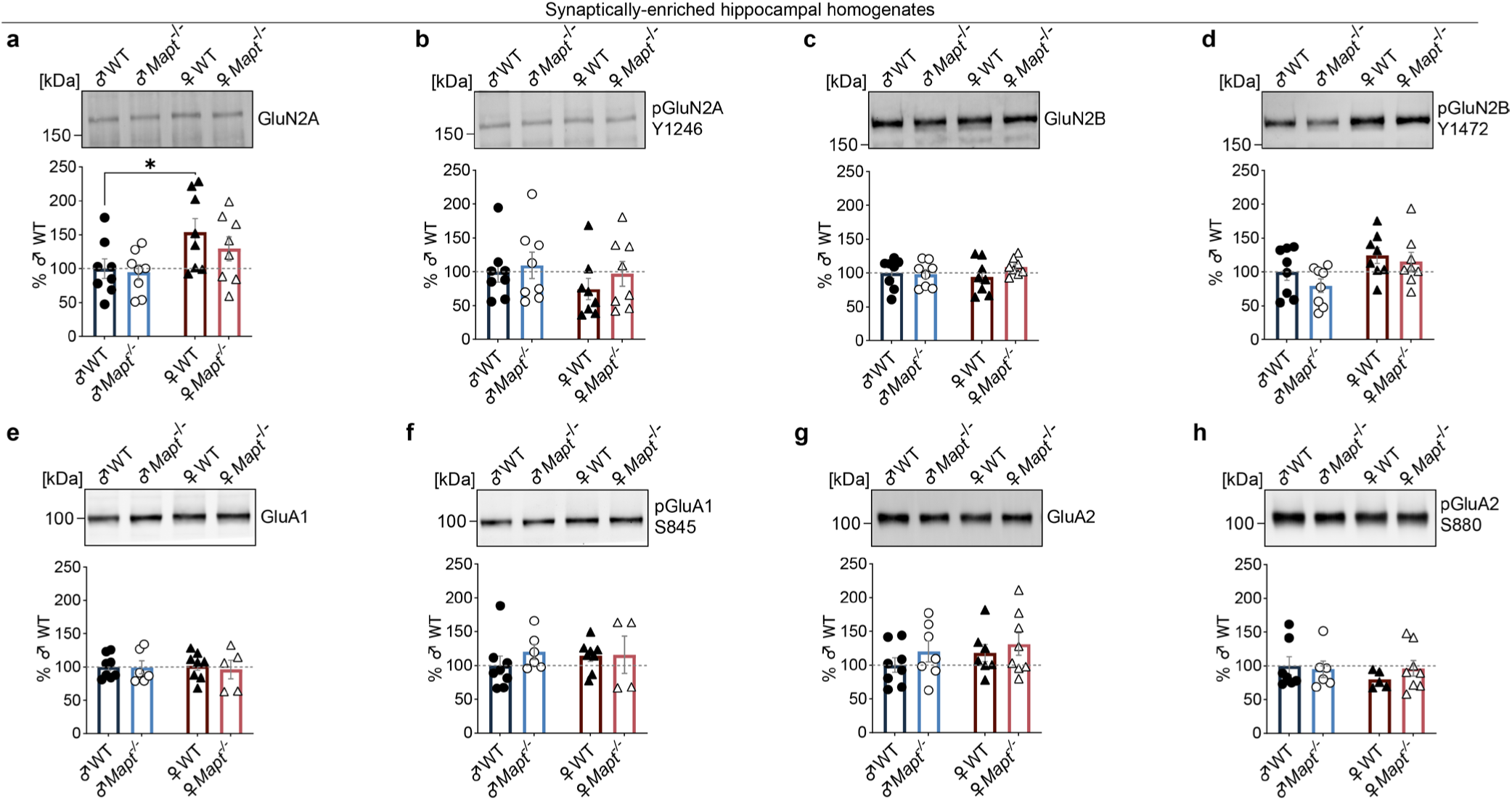
Synaptic protein and phosphorylation levels of NMDAR and AMPAR subunits are not impacted by tau deletion in either sex. **a-h**, Representative immunoblots (upper) and quantification (lower) for protein and phosphorylation levels of NMDAR (a-d) and AMPAR (e-h) subunits in synaptically-enriched rat hippocampal homogenates. There was a main effect for sex (two-way ANOVA: *P* = 0.01) on synaptic GluN2A levels such that female WTs (154.1 ± 20.0%) expressed more GluN2A relative to male WTs (100.0 ± 14.4%; Šídák’s post hoc test: *P* = 0.02) (a). A main effect of sex on GluN2B-Y1472 phosphorylation (two-way ANOVA: *P* = 0.02) was not significant between sexes of WT rats (Šídák’s post hoc test: *P* = 0.3) (d). Subunit levels are normalized to total (stain-free) protein levels, while phosphorylation epitopes are normalized to their respective subunit protein levels. All data are expressed as % of male WT (*n* = 4-8 per group). Comparisons are by two-way ANOVA followed by Šídák’s post hoc test. Detailed statistics are shown in Supplementary Table 3. \**P* < 0.05.

**Extended Data Figure 5.**
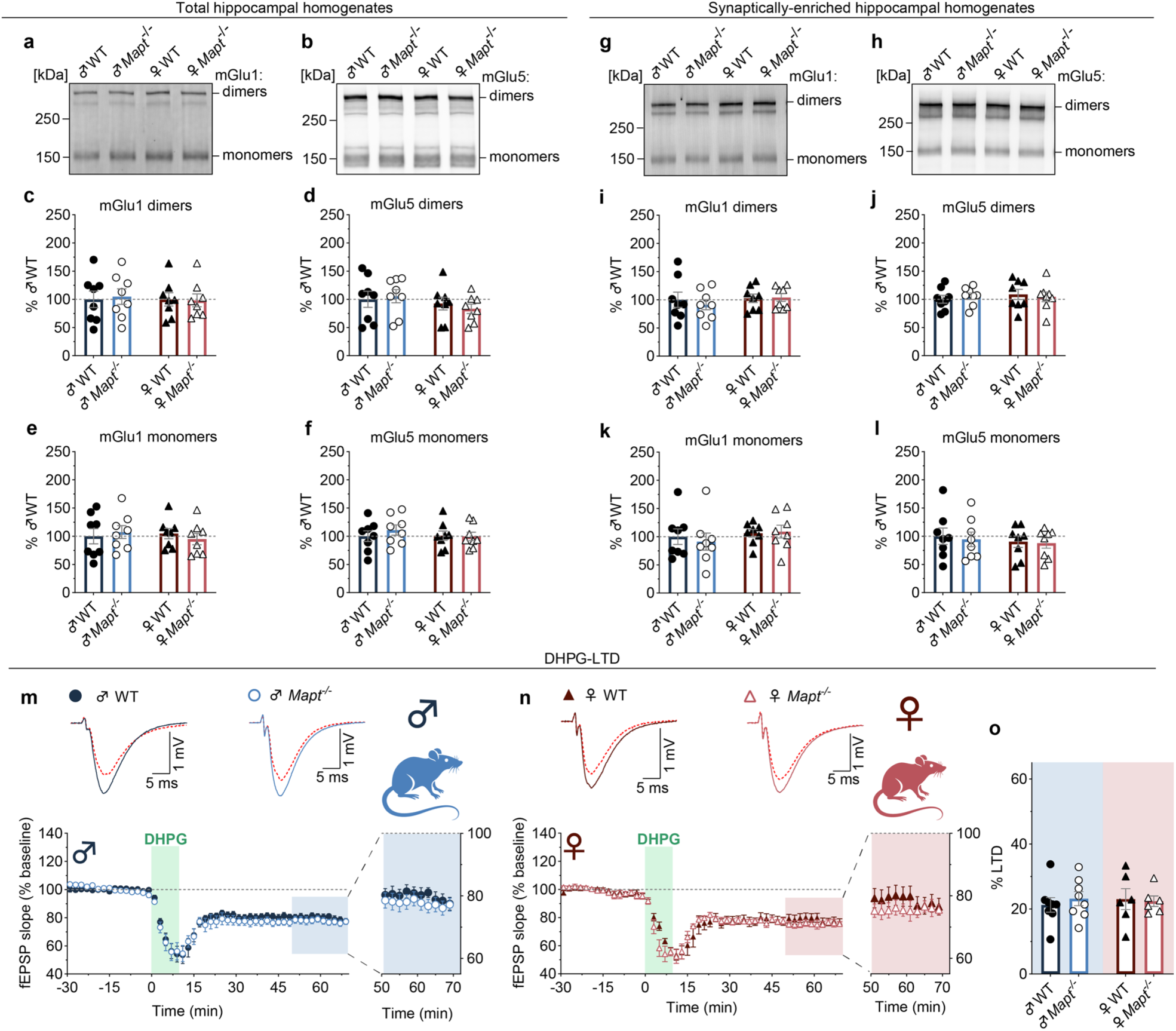
Similar expression of mGluR_I_ subtypes and levels of DHPG-LTD in both sexes of WT and *Mapt^-/-^* rats. **a**-**l**, Representative blot images and quantification of mGu1 and mGlu5 monomer and dimer levels in total (**a**-**f**) and synaptically enriched (**g**-**l**) hippocampal homogenates from both sexes of WT and *Mapt^-^*^/-^ rats. There were no genotype or sex differences in total or synaptic expression levels of mGlu1 or mGlu5 (*n* = 8 per group). mGlu1 and mGlu5 protein levels are normalized to total (stain free) protein levels and expressed as % of male WT. **m**-**o**, fEPSP recordings showing levels of LTD induced by a 10 min application of the mGluR_I_ agonist (*S*)-3,5-DHPG (50 µM) in juvenile male (**m**; *n* = 8 per genotype) and female (**n**; *n* = 6 per genotype) WT and *Mapt^-/-^* rats. DHPG-LTD levels were similar in WT and *Mapt^-/-^* rats of both sexes (**o**). Expanded regions of the time plots show the final 20 min of recordings. Representative traces show superimposed fEPSPs from the baseline period (solid line) and the end of the recording session (dashed line). Levels of LTD were calculated as the percent change from baseline in the last 6 min of recordings. Comparisons were made using two-way ANOVA. Detailed statistics are shown in Supplementary Tables 2 and 3.

**Extended Data Figure 6.**
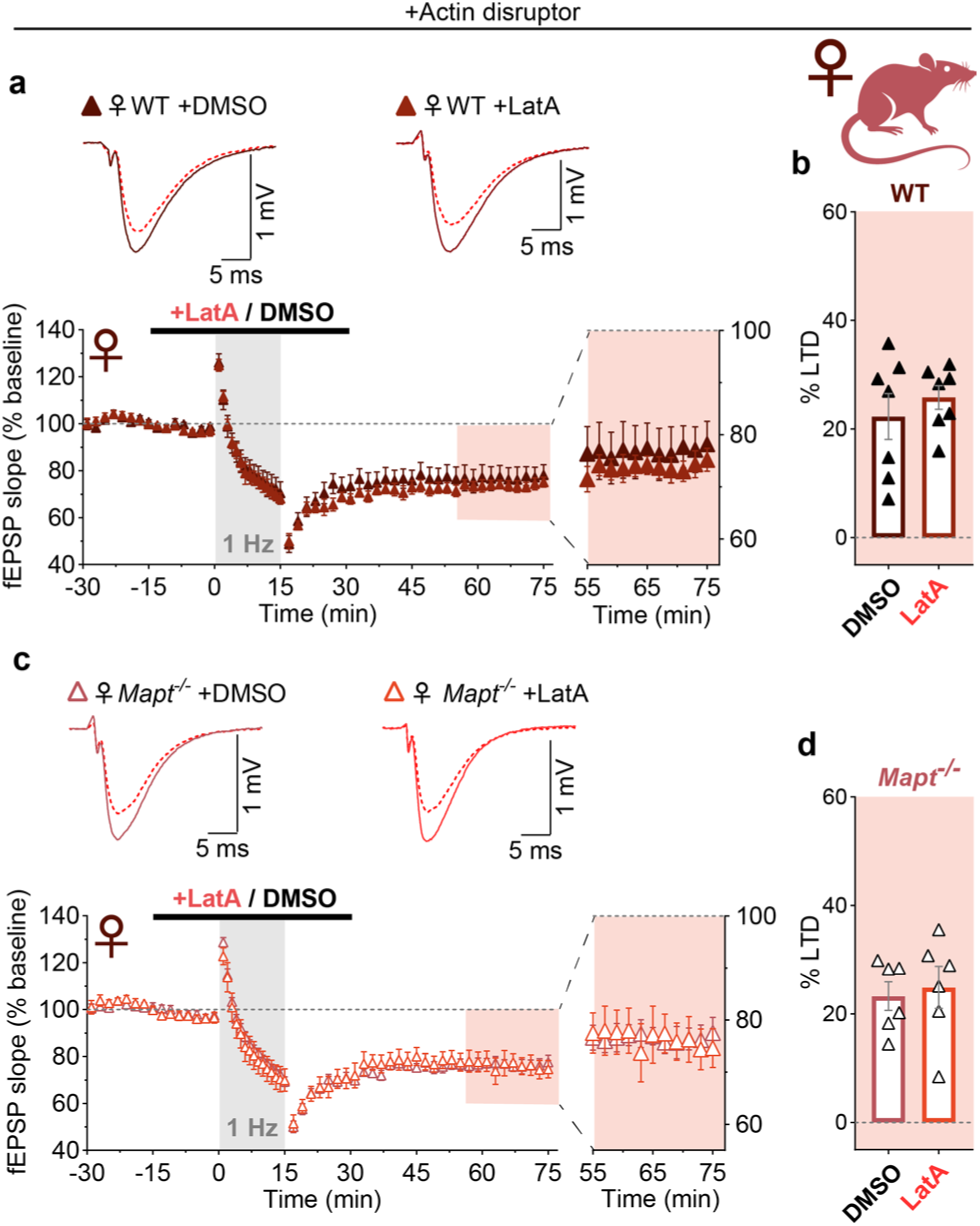
LFS-LTD is not affected by latrunculin-A in female WT or *Mapt^-/-^* rats. **a**-**d**, fEPSP recordings examining the impact of latrunculin A (LatA; 0.1 µM) on LFS-LTD in slices from female WT and *Mapt^-/-^* rats. LatA or DMSO (0.001%) was applied for 15 min prior to, during, and 15 min post LFS induction. In female WT rats (**a**,**b**) LatA treatment had no effect on LTD magnitude (25.8 ± 2.2%) relative to DMSO-treated slices (22.3 ± 4.2%, *n* = 7 per genotype; *P* = 0.5). Similarly, LTD levels in female *Mapt^-/-^* rats (**c**,**d**) were not significantly impacted by LatA treatment (24.9 ± 3.9%) relative to LTD in DMSO controls (23.3 ± 2.6%, *n* = 6 per genotype; *P* = 0.7). The final 20 min of recordings are shown in expanded regions. Representative traces show superimposed fEPSPs from the baseline period (solid line) and the end of the recording session (dashed line). LTD levels were calculated as the percent change from baseline in the last 6 min of recordings. Comparisons were made using unpaired two-tailed Student’s *t*-tests. Detailed statistics are shown in Supplementary Table 3.

## Notes

### Competing Interest Statement

The authors have declared no competing interest.

## REFERENCES

1. Goedert, M., Crowther, R. A., Scheres, S. H. W. & Spillantini, M. G. Tau and neurodegeneration. Cytoskeleton at 10.1002/cm.21812 (2024).

2. Tzioras, M., McGeachan, R. I., Durrant, C. S. & Spires-Jones, T. L. Synaptic degeneration in Alzheimer disease. Nature Reviews Neurology at 10.1038/s41582-022-00749-z (2023).

3. Parra Bravo, C., Naguib, S. A. & Gan, L. Cellular and pathological functions of tau. Nat. Rev. Mol. Cell Biol. 2024 2511 25, 845–864 (2024).

4. Pichet Binette, A., et al. Proteomic changes in Alzheimer’s disease associated with progressive Aβ plaque and tau tangle pathologies. Nat. Neurosci. 2024 2710 27, 1880–1891 (2024).

5. Frandemiche, M. L. et al. Activity-dependent tau protein translocation to excitatory synapse is disrupted by exposure to amyloid-beta oligomers. J. Neurosci. 34, 6084–6097 (2014).

6. Hoover, B. R. et al. Tau Mislocalization to Dendritic Spines Mediates Synaptic Dysfunction Independently of Neurodegeneration. Neuron (2010) doi:10.1016/j.neuron.2010.11.030.

7. Ittner, L. M. et al. Dendritic function of tau mediates amyloid-beta toxicity in Alzheimer’s disease mouse models. Cell 142, 387–397 (2010).

8. Kimura, T. et al. Microtubule-associated protein tau is essential for long-term depression in the hippocampus. Philos. Trans. R. Soc. B Biol. Sci. 369, 20130144 (2014).

9. Xia, D., Gutmann, J. M. & Gotz, J. Mobility and subcellular localization of endogenous, gene-edited Tau differs from that of over-expressed human wild-type and P301L mutant Tau. Sci. Rep. (2016) doi:10.1038/srep29074.

10. Robbins, M., Clayton, E. & Kaminski Schierle, G. S. Synaptic tau: A pathological or physiological phenomenon? Acta Neuropathologica Communications at 10.1186/s40478-021-01246-y (2021).

11. Regan, P. et al. Tau phosphorylation at serine 396 residue is required for hippocampal LTD. J. Neurosci. 35, 4804–4812 (2015).

12. Biundo, F., Del Prete, D., Zhang, H., Arancio, O. & D’Adamio, L. A role for tau in learning, memory and synaptic plasticity. Sci. Rep. 8, 3184 (2018).

13. Ahmed, T. et al. Cognition and hippocampal synaptic plasticity in mice with a homozygous tau deletion. Neurobiol. Aging 35, 2474–2478 (2014).

14. Watamura, N. et al. In vivo hyperphosphorylation of tau is associated with synaptic loss and behavioral abnormalities in the absence of tau seeds. Nat. Neurosci. 2024 282 28, 293–307 (2024).

15. Taddei, R. N. & Duff, K. E. Synapse vulnerability and resilience across the clinical spectrum of dementias. Nat. Rev. Neurol. 2025 217 21, 353–369 (2025).

16. Collingridge, G. L., Peineau, S., Howland, J. G. & Wang, Y. T. Long-term depression in the CNS. Nature Reviews Neuroscience at 10.1038/nrn2867 (2010).

17. Dudek, S. M. & Bear, M. F. Homosynaptic long-term depression in area CA1 of hippocampus and effects of N-methyl-D-aspartate receptor blockade. Proc. Natl. Acad. Sci. U. S. A. (1992) doi:10.1073/pnas.89.10.4363.

18. Dejanovic, B., Sheng, M. & Hanson, J. E. Targeting synapse function and loss for treatment of neurodegenerative diseases. Nat. Rev. Drug Discov. 23, 23–42 (2024).

19. Bliss, T. V. P. & Collingridge, G. L. A synaptic model of memory: Long-term potentiation in the hippocampus. Nature at 10.1038/361031a0 (1993).

20. Ayers, J. et al. Severe neurodegeneration in brains of transgenic rats producing human tau prions. Acta Neuropathol. 148, (2024).

21. Do Carmo, S. & Cuello, A. C. Modeling Alzheimer’s disease in transgenic rats. Molecular Neurodegeneration at 10.1186/1750-1326-8-37 (2013).

22. Gertsenstein, M. & Nutter, L. M. J. Production of knockout mouse lines with Cas9. Methods (2021) doi:10.1016/j.ymeth.2021.01.005.

23. Gallego-Rudolf, J., Wiesman, A. I., Pichet Binette, A., Villeneuve, S. & Baillet, S. Synergistic association of Aβ and tau pathology with cortical neurophysiology and cognitive decline in asymptomatic older adults. Nat. Neurosci. 2024 2711 27, 2130–2137 (2024).

24. Tracy, T. E. et al. Tau interactome maps synaptic and mitochondrial processes associated with neurodegeneration. Cell (2022) 10.1016/j.cell.2021.12.041.

25. Hill, E., Karikari, T. K., Moffat, K. G., Richardson, M. J. E. & Wall, M. J. Introduction of Tau oligomers into cortical neurons alters action potential dynamics and disrupts synaptic transmission and plasticity. eNeuro 6, (2019).

26. Regan, P. et al. Tau phosphorylation at serine 396 residue is required for hippocampal LTD. J. Neurosci. (2015) doi:10.1523/JNEUROSCI.2842-14.2015.

27. Maggio, N. & Segal, M. Persistent changes in ability to express long-term potentiation/depression in the rat hippocampus after juvenile/adult stress. Biol. Psychiatry (2011) doi:10.1016/j.biopsych.2010.11.026.

28. Mulkey, R. M. & Malenka, R. C. Mechanisms underlying induction of homosynaptic long-term depression in area CA1 of the hippocampus. Neuron (1992) doi:10.1016/0896-6273(92)90248-C.

29. Volk, L. J., Daly, C. A. & Huber, K. M. Differential roles for group 1 mGluR subtypes in induction and expression of chemically induced hippocampal long-term depression. J. Neurophysiol. (2006) doi:10.1152/jn.00383.2005.

30. Oliet, S. H. R., Malenka, R. C. & Nicoll, R. A. Two distinct forms of long-term depression coexist in CA1 hippocampal pyramidal cells. Neuron (1997) doi:10.1016/S0896-6273(00)80336-0.

31. Bashir, Z. I., Jane, D. E., Sunter, D. C., Watkins, J. C. & Collingridge, G. L. Metabotropic glutamate receptors contribute to the induction of long-term depression in the CA1 region of the hippocampus. Eur. J. Pharmacol. (1993) doi:10.1016/0014-2999(93)91009-C.

32. Palmer, M. J., Irving, A. J., Seabrook, G. R., Jane, D. E. & Collingridge, G. L. The group I mGlu receptor agonist DHPG induces a novel form of LTD in the CA1 region of the hippocampus. Neuropharmacology (1997) doi:10.1016/S0028-3908(97)00181-0.

33. Ralph, L. T., John, G., Collingridge, G. L. & Tidball, P. Sex-dependence of synaptic depression induced by activation of metabotropic glutamate receptors in rat hippocampus. Brain Neurosci. Adv. 8, 23982128231223580 (2024).

34. Scheefhals, N., Westra, M. & MacGillavry, H. D. mGluR5 is transiently confined in perisynaptic nanodomains to shape synaptic function. Nat. Commun. (2023) doi:10.1038/s41467-022-35680-w.

35. Farr, C. D. et al. Proteomic analysis of native metabotropic glutamate receptor 5 protein complexes reveals novel molecular constituents. J. Neurochem. (2004) doi:10.1111/j.1471-4159.2004.02735.x.

36. Sergé, A., Fourgeaud, L., Hémar, A. & Choquet, D. Active surface transport of metabotropic glutamate receptors through binding to microtubules and actin flow. J. Cell Sci. (2003) doi:10.1242/jcs.00822.

37. Cabrales Fontela, Y., et al. Multivalent cross-linking of actin filaments and microtubules through the microtubule-associated protein Tau. Nat Commun 8, 1981 (2017).

38. Fujiwara, I., Zweifel, M. E., Courtemanche, N. & Pollard, T. D. Latrunculin A Accelerates Actin Filament Depolymerization in Addition to Sequestering Actin Monomers. Curr. Biol. (2018) doi:10.1016/j.cub.2018.07.082.

39. Chen, Y., Bourne, J., Pieribone, V. A. & Fitzsimonds, R. M. The role of actin in the regulation of dendritic spine morphology and bidirectional synaptic plasticity. Neuroreport (2004) doi:10.1097/00001756-200404090-00018.

40. Zhou, Z., Hu, J., Passafaro, M., Xie, W. & Jia, Z. GluA2 (GluR2) regulates metabotropic glutamate receptor-dependent long-term depression through N-cadherin-dependent and cofilin-mediated actin reorganization. J. Neurosci. (2011) doi:10.1523/JNEUROSCI.3869-10.2011.

41. Okamoto, K. I., Nagai, T., Miyawaki, A. & Hayashi, Y. Rapid and persistent modulation of actin dynamics regulates postsynaptic reorganization underlying bidirectional plasticity. Nat. Neurosci. (2004) doi:10.1038/nn1311.

42. Ku, H. Y., Huang, Y. F., Chao, P. H., Huang, C. C. & Hsu, K. Sen. Neonatal isolation delays the developmental decline of long-term depression in the CA1 region of rat hippocampus. Neuropsychopharmacology (2008) doi:10.1038/npp.2008.36.

43. McKean, N. E., Handley, R. R. & Snell, R. G. A review of the current mammalian models of alzheimer’s disease and challenges that need to be overcome. Int. J. Mol. Sci. (2021) doi:10.3390/ijms222313168.

44. Jaramillo, S., Zador, A. M. & Jaramillo, S. Mice and rats achieve similar levels of performance in an adaptive decision-making task. Front. Syst. Neurosci. (2014) doi:10.3389/fnsys.2014.00173.

45. Cohen, R. M. et al. A transgenic alzheimer rat with plaques, tau pathology, behavioral impairment, oligomeric Aβ, and frank neuronal loss. J. Neurosci. (2013) doi:10.1523/JNEUROSCI.3672-12.2013.

46. Ellenbroek, B. & Youn, J. Rodent models in neuroscience research: Is it a rat race? DMM Dis. Model. Mech. (2016) doi:10.1242/dmm.026120.

47. Hanes, J. et al. Rat tau proteome consists of six tau isoforms: Implication for animal models of human tauopathies. J. Neurochem. (2009) doi:10.1111/j.1471-4159.2009.05869.x.

48. Takuma, H., Arawaka, S. & Mori, H. Isoforms changes of tau protein during development in various species. Dev. Brain Res. (2003) doi:10.1016/S0165-3806(03)00056-7.

49. Tuerde, D. et al. Isoform-independent and-dependent phosphorylation of microtubule-associated protein tau in mouse brain during postnatal development. J. Biol. Chem. (2018) doi:10.1074/jbc.M117.798918.

50. McMillan, P. et al. Tau isoform regulation is region-and cell-specific in mouse brain. J. Comp. Neurol. (2008) doi:10.1002/cne.21867.

51. He, Z. et al. Transmission of tauopathy strains is independent of their isoform composition. Nat. Commun. (2020) doi:10.1038/s41467-019-13787-x.

52. Yang, Y., Ondrejcak, T., Hu, N. W., Klyubin, I. & Rowan, M. J. Divergent disruptive effects of soluble recombinant tau assemblies on synaptic plasticity in vivo. Mol. Brain 18, 1–14 (2025).

53. Hu, N. W. et al. Patient-derived tau and amyloid-β facilitate long-term depression in vivo: role of tumour necrosis factor-α and the integrated stress response. Brain Commun. 6, fcae333 (2024).

54. Harada, A. et al. Altered microtubule organization in small-calibre axons of mice lacking tau protein. Nature (1994) doi:10.1038/369488a0.

55. Buchholz, S. & Zempel, H. The six brain-specific TAU isoforms and their role in Alzheimer’s disease and related neurodegenerative dementia syndromes. Alzheimer’s Dement. 20, 3606 (2024).

56. Lopes, S. et al. Tau protein is essential for stress-induced brain pathology. Proc. Natl. Acad. Sci. U. S. A. (2016) doi:10.1073/pnas.1600953113.

57. Francis, C. et al. Divergence of RNA localization between rat and mouse neurons reveals the potential for rapid brain evolution. BMC Genomics (2014) doi:10.1186/1471-2164-15-883.

58. Hagena, H. & Manahan-Vaughan, D. Interplay of hippocampal long-term potentiation and long-term depression in enabling memory representations. Philos. Trans. B 379, (2024).

59. Belkacemi, K., Rondard, P., Pin, J. P. & Prézeau, L. Heterodimers Revolutionize the Field of Metabotropic Glutamate Receptors. Neuroscience (2024) doi:10.1016/J.NEUROSCIENCE.2024.06.013.

60. Moult, P. R., Corrêa, S. A. L., Collingridge, G. L., Fitzjohn, S. M. & Bashir, Z. I. Co-activation of p38 mitogen-activated protein kinase and protein tyrosine phosphatase underlies metabotropic glutamate receptor-dependent long-term depression. J. Physiol. (2008) doi:10.1113/jphysiol.2008.153122.

61. Huber K M, Kayser M S & Bear M F. Role for rapid dendritic protein synthesis in hippocampal mGluR-dependent long-term depression. Science (80-.). (2000).

62. He, H. J. et al. The proline-rich domain of tau plays a role in interactions with actin. BMC Cell Biol. (2009) doi:10.1186/1471-2121-10-81.

63. Gall, C. M., Le, A. A. & Lynch, G. Sex differences in synaptic plasticity underlying learning. Journal of Neuroscience Research at 10.1002/jnr.24844 (2023).

64. Le, A. A. et al. Prepubescent female rodents have enhanced hippocampal LTP and learning relative to males, reversing in adulthood as inhibition increases. Nat. Neurosci. (2022) doi:10.1038/s41593-021-01001-5.

65. Le, A. A. et al. Metabotropic NMDA Receptor Signaling Contributes to Sex Differences in Synaptic Plasticity and Episodic Memory. (2024) doi:10.1101/2024.01.26.577478.

66. Ondrejcak, T. et al. Rapidly reversible persistent long-term potentiation inhibition by patient-derived brain tau and amyloid ß proteins. Philos. Trans. R. Soc. Lond. B. Biol. Sci. 379, (2024).

67. Shipton, O. A. et al. Tau protein is required for amyloid beta-induced impairment of hippocampal long-term potentiation. J. Neurosci. 31, 1688–1692 (2011).

68. Jo, J. et al. Aβ1-42 inhibition of LTP is mediated by a signaling pathway involving caspase-3, Akt1 and GSK-3β. Nat. Neurosci. (2011) doi:10.1038/nn.2785.

69. Ng, A. N., Salter, E. W., Georgiou, J., Bortolotto, Z. A. & Collingridge, G. L. Amyloid-β1-42 oligomers enhance mGlu5R-dependent synaptic weakening via NMDAR activation and complement C5aR1 signaling. iScience (2023) doi:10.1016/j.isci.2023.108412.

70. Um, J. W. et al. Metabotropic Glutamate Receptor 5 Is a Coreceptor for Alzheimer Aβ Oligomer Bound to Cellular Prion Protein. Neuron (2013) doi:10.1016/j.neuron.2013.06.036.

71. Hu, N. W. et al. MGlu5 receptors and cellular prion protein mediate amyloid-β-facilitated synaptic long-term depression in vivo. Nat. Commun. (2014) doi:10.1038/ncomms4374.

72. Abd-Elrahman, K. S. et al. Aβ oligomers induce pathophysiological mGluR5 signaling in Alzheimer’s disease model mice in a sex-selective manner. Sci. Signal. (2020) doi:10.1126/scisignal.abd2494.

73. Wang, Y. et al. Tau pathology is associated with postsynaptic metabotropic glutamate receptor 5 (mGluR5) in early Alzheimer’s disease in a sex-specific manner. Alzheimer’s Dement. (2025) 10.1002/alz.70004.

74. Chang, C. W., Shao, E. & Mucke, L. Tau: Enabler of diverse brain disorders and target of rapidly evolving therapeutic strategies. Science at 10.1126/science.abb8255 (2021).

75. Castro-Aldrete, L. et al. Alzheimer disease seen through the lens of sex and gender. Nat. Rev. Neurol. (2025) doi:10.1038/S41582-025-01071-0,.

76. Congdon, E. E., Ji, C., Tetlow, A. M., Jiang, Y. & Sigurdsson, E. M. Tau-targeting therapies for Alzheimer disease: current status and future directions. Nature Reviews Neurology at 10.1038/s41582-023-00883-2 (2023).

77. Filipiak, W. E. & Saunders, T. L. Advances in transgenic rat production. Transgenic Res. (2006) doi:10.1007/s11248-006-9002-x.

78. Gertsenstein, M. & Nutter, L. M. J. Engineering Point Mutant and Epitope-Tagged Alleles in Mice Using Cas9 RNA-Guided Nuclease. Curr. Protoc. Mouse Biol. (2018) doi:10.1002/cpmo.40.

79. Gonçalves, R. A., Wijesekara, N., Fraser, P. E. & De Felice, F. G. Behavioral Abnormalities in Knockout and Humanized Tau Mice. Front. Endocrinol. (Lausanne). (2020) doi:10.3389/fendo.2020.00124.

80. Truett, G. E. et al. Preparation of PCR-quality mouse genomic dna with hot sodium hydroxide and tris (HotSHOT). Biotechniques (2000) doi:10.2144/00291bm09.

81. Anderson, W. W. & Collingridge, G. L. Capabilities of the WinLTP data acquisition program extending beyond basic LTP experimental functions. J. Neurosci. Methods (2007) doi:10.1016/j.jneumeth.2006.12.018.

82. Hallett, P. J., Collins, T. L., Standaert, D. G. & Dunah, A. W. Biochemical fractionation of brain tissue for studies of receptor distribution and trafficking. Curr. Protoc. Neurosci. (2008) doi:10.1002/0471142301.ns0116s42.

