## Supplementary Information for "Tau deletion results in sex-dependent modulation of synaptic weakening in rat hippocampus"

Supplementary Information for **BIORXIV/2025/665155**. Ralph et al., 2025:
**Tau deletion results in sex-dependent modulation of synaptic weakening in rat hippocampus**

**Supplementary Table 1**. Statistics summary for WT and *Mapt*$^{-/-}$ rat body, brain, and hippocampus mass measurements.

**Supplementary Table 2**. Statistics summary for WT and *Mapt*$^{-/-}$ rat and mouse electrophysiology.

**Supplementary Table 3**. Statistics summary for WT and *Mapt*$^{-/-}$ rat immunoblotting.

**Supplementary Figure 1**. Lane order, full-blot images, total protein images, and primary antibody targets for representative immunoblots for *Mapt*$^{-/-}$ rat immunoblotting experiments.

**Supplementary Methods Table 1**. Antibodies used for *Mapt*$^{-/-}$ rat immunoblotting experiments.

---

**Supplementary Table 1**. Statistics summary for WT and *Mapt*$^{-/-}$ rat body, brain and hippocampus mass measurements.

| Figure number and experiment: | Group (number of rodents), and mean ± SEM: | Statistical tests, *F/t*-statistic, degrees of freedoms (dfn,d), and *P*-values | Post-hoc test and *P*-values: |
|---|---|---|---|
| **Fig. 1e.** Body mass | Body mass (g):<br>**Male WT rats (7)**: 43.5 ± 4.2<br>**Male *Mapt*$^{-/-}$ rats (7)**: 36.5 ± 2.4<br>**Female WT rats (9)**: 38.9 ± 2.7<br>**Female *Mapt*$^{-/-}$ rats (9)**: 37.3 ± 4.7 | Two-way ANOVA:<br>**Interaction**: $F_{(1, 28)}$ = 0.5, $P$ = 0.5<br>**Sex effect**: $F_{(1, 28)}$ = 0.3, $P$ = 0.6<br>**Genotype effect**: $F_{(1, 28)}$ = 1.3, $P$ = 0.3 | N/A (no significant interactions, or main effects in two-way ANOVA) |
| **Fig. 1e.** Brain mass | Brain mass (g):<br>**Male WT rats (4)**: 14.8 ± 0.5<br>**Male *Mapt*$^{-/-}$ rats (4)**: 13.4 ± 0.4<br>**Female WT rats (8)**: 13.9 ± 0.6<br>**Female *Mapt*$^{-/-}$ rats (8)**: 13.0 ± 0.5 | Two-way ANOVA:<br>**Interaction**: $F_{(1, 20)}$ = 0.2, $P$ = 0.7<br>**Sex effect**: $F_{(1, 20)}$ = 1.1, $P$ = 0.3<br>**Genotype effect**: $F_{(1, 20)}$ = 3.6, $P$ = 0.07 | N/A (no significant interactions, or main effects in two-way ANOVA) |
| **Fig. 1e.** Hippocampus mass | Hippocampus mass (mg):<br>**Male WT rats (4)**: 122.9 ± 4.5<br>**Male *Mapt*$^{-/-}$ rats (4)**: 117.6 ± 3.0<br>**Female WT rats (8)**: 116.8 ± 6.4<br>**Female *Mapt*$^{-/-}$ rats (8)**: 107.4 ± 9.7 | Two-way ANOVA:<br>**Interaction**: $F_{(1, 20)}$ = 0.06, $P$ = 0.8<br>**Sex effect**: $F_{(1, 20)}$ = 0.9, $P$ = 0.4<br>**Genotype effect**: $F_{(1, 20)}$ = 0.7, $P$ = 0.4 | N/A (no significant interactions, or main effects in two-way ANOVA) |

**Supplementary Table 2**. Statistics summary for WT and *Mapt*$^{-/-}$ rat and mouse electrophysiology.

| Figure number and experiment: | Group (number of rodents) and mean ± SEM: | Statistical tests, *F/t*-statistic, degrees of freedoms (dfn,d), and *P*-values | Post-hoc test and *P*-values: |
|---|---|---|---|
| **Fig. 1f-h**. LTD levels in male and female WT versus *Mapt*$^{-/-}$ rats (dataset #1) | % LTD:<br>**Male WT rats (8)**: 17.9 ± 2.7<br>**Male *Mapt*$^{-/-}$ rats (8)**: 31.8 ± 3.1<br>**Female WT rats (8)**: 12.6 ± 2.4<br>**Female *Mapt*$^{-/-}$ rats (8)**: 15.3 ± 1.5 | Two-way ANOVA:<br>**Interaction**: $F_{(1, 28)}$ = 5.09, *P* = 0.03<br>**Sex effect**: $F_{(1, 28)}$ = 19.2, *P* = 0.0002<br>**Genotype effect**: $F_{(1, 28)}$ = 11.2, *P* = 0.002 | Selected Šídák's post hoc test (adj. *P*-values):<br>**Male WT vs. *Mapt*$^{-/-}$**: *P* = 0.0014<br>**Female WT vs. *Mapt*$^{-/-}$**: *P* = 0.8314<br>**WT male vs. female**: *P* = 0.8514 |
| **Fig. 1i and Extended Data Fig. 3a**. Axon activation male and female WT versus *Mapt*$^{-/-}$ rats | Slope of FV amplitude vs. stimulation intensity (mV/μA):<br>**Male WT rats (10)**: -0.00861 ± 0.00092<br>**Male *Mapt*$^{-/-}$ rats (10)**: -0.00814 ± 0.00064<br>**Female WT rats (10)**: -0.00860 ± 0.00064<br>**Female *Mapt*$^{-/-}$ rats (10)**: -0.00875 ± 0.00085 | Two-way ANOVA:<br>**Interaction**: $F_{(1, 36)}$ = 0.00032, *P* = 0.9<br>**Sex effect**: $F_{(1, 36)}$ = 0.12, *P* = 0.7<br>**Genotype effect**: $F_{(1, 36)}$ = 1.88, *P* = 0.28 | N/A (no significant interactions, or main effects in two-way ANOVA) |
| **Fig. 1j and Extended Data Fig. 3b**. Basal synaptic transmission male and female WT versus and *Mapt*$^{-/-}$ rats | Slope of fEPSP slope vs. FV amplitude (mV/ms·mV$^{-1}$):<br>**Male WT rats (10)**: 2.0 ± 0.1<br>**Male *Mapt*$^{-/-}$ rats (10)**: 2.3 ± 0.2<br>**Female WT rats (10)**: 1.8 ± 0.2<br>**Female *Mapt*$^{-/-}$ rats (10)**: 2.0 ± 0.2 | Two-way ANOVA:<br>**Interaction**: $F_{(1, 36)}$ = 0.0043, *P* = 0.95<br>**Sex effect**: $F_{(1, 36)}$ = 2.27, *P* = 0.14<br>**Genotype effect**: $F_{(1, 36)}$ = 2.3, *P* = 0.14 | N/A (no significant interactions, or main effects in two-way ANOVA) |
| **Fig. 1k and Extended Data Fig. 3c**. Paired-pulse facilitation male and female WT versus *Mapt*$^{-/-}$ rats | Paired-pulse ratio (fEPSP2/1 – 50 ms interval):<br>**Male WT rats (10)**: 2.0 ± 0.1<br>**Male *Mapt*$^{-/-}$ rats (10)**: 2.0 ± 0.1<br>**Female WT rats (10)**: 1.9 ± 0.1<br>**Female *Mapt*$^{-/-}$ rats (10)**: 2.1 ± 0.1 | Two-way ANOVA:<br>**Interaction**: $F_{(1, 36)}$ = 1.4, *P* = 0.24<br>**Sex effect**: $F_{(1, 36)}$ = 0.3, *P* = 0.59<br>**Genotype effect**: $F_{(1, 36)}$ = 1.17, *P* = 0.29 | N/A (no significant interactions, or main effects in two-way ANOVA) |
| **Fig. 2a,b**. LTD levels in male WT versus *Mapt*$^{-/-}$ mice | % LTD:<br>**Male WT mice (6)**: 26.4 ± 1.6<br>**Male *Mapt*$^{-/-}$ mice (6)**: 14.3 ± 3.5 | Unpaired Student's *t*-test:<br>***t*-statistic**: 3.16<br>**Df**: 10<br>***P*-value**: 0.0102 | N/A |
| **Fig. 2c,d**. LTD levels in male WT versus *Mapt*$^{-/-}$ rats (dataset #2) | % LTD:<br>**Male WT rats (6)**: 22.0 ± 4.3<br>**Male *Mapt*$^{-/-}$ rats (6)**: 40.3 ± 3.1 | Unpaired Student's *t*-test:<br>***t*-statistic**: 3.44<br>**Df**: 10<br>***P*-value**: 0.0063 | N/A |
| **Fig. 2e,f**. LTD levels in female WT versus *Mapt*$^{-/-}$ mice | % LTD:<br>**Female WT mice (6)**: 15.2 ± 1.6<br>**Female *Mapt*$^{-/-}$ mice (6)**: 12.9 ± 4.7 | Unpaired Student's *t*-test:<br>***t*-statistic**: 0.95<br>**Df**: 10<br>***P*-value**: 0.36 | N/A |
| **Fig. 2g,h**. LTD levels in female WT versus *Mapt*$^{-/-}$ rat (dataset #2) | % LTD:<br>**Female WT rats (6)**: 19.3 ± 3.7<br>**Female *Mapt*$^{-/-}$ rats (6)**: 28.1 ± 2.3 | Unpaired Student's *t*-test:<br>***t*-statistic**: 2.0<br>**Df**: 10<br>***P*-value**: 0.07 | N/A |
| **Fig. 3a-c**. LTD+D-AP5 levels in male and female WT versus *Mapt*$^{-/-}$ rats (dataset #1) | % LTD:<br>**Male WT rats (8)**: 0.1 ± 2.5<br>**Male *Mapt*$^{-/-}$ rats (8)**: 12.7 ± 1.9<br>**Female WT rats (8)**: 0.8 ± 2.6<br>**Female *Mapt*$^{-/-}$ rats (8)**: 0.9 ± 3.1 | Two-way ANOVA:<br>**Interaction**: $F_{(1, 28)}$ = 5.9, *P* = 0.02<br>**Sex effect**: $F_{(1, 28)}$ = 4.7, *P* = 0.04<br>**Genotype effect**: $F_{(1, 28)}$ = 6.07, *P* = 0.02 | Selected Šídák's post hoc test (adj. *P*-values):<br>**Male WT vs. *Mapt*$^{-/-}$**: *P* = 0.005<br>**Female WT vs. *Mapt*$^{-/-}$**: *P* > 0.99<br>**WT male vs. female**: *P* > 0.99 |
| **Fig. 4a-c**. LTD+AP5 (dataset #2) and LTD+AP5+YM+MTEP in male WT versus *Mapt*$^{-/-}$ rats | % LTD:<br>**Male WT rats +AP5 (8)**: 1.9 ± 2.5<br>**Male *Mapt*$^{-/-}$ rats +AP5 (8)**: 10.5 ± 2.0<br>**Male WT rats +AP5+YM+MTEP (8)**: 0.7 ± 1.2<br>**Male *Mapt*$^{-/-}$ rats +AP5+YM+MTEP (8)**: -1.7 ± 2.1 | Two-way ANOVA:<br>**Interaction**: $F_{(1, 28)}$ = 7.5, *P* = 0.01<br>**YM+MTEP effect**: $F_{(1, 28)}$ = 11.04, *P* = 0.003<br>**Genotype effect**: $F_{(1, 28)}$ = 2.4, *P* = 0.14 | Selected Šídák's post hoc test (adj. *P*-values):<br>**D-AP5: WT vs. *Mapt*$^{-/-}$**: *P* = 0.021<br>**D-AP5+YM+MTEP: WT vs. *Mapt*$^{-/-}$**: *P* = 0.88<br>**WT: D-AP5 vs. D-AP5+YM+MTEP**: *P* = 0.99<br>***Mapt*$^{-/-}$: D-AP5 vs. D-AP5+YM+MTEP**: *P* = 0.0008 |
| **Fig. 4d-f**. LTD (dataset #3) and LTD+YM+MTEP in male WT versus *Mapt*$^{-/-}$ rats | % LTD:<br>**Male WT rats (10)**: 22.4 ± 2.6<br>**Male *Mapt*$^{-/-}$ rats (10)**: 33.8 ± 2.9<br>**Male WT rats +YM+MTEP (10)**: 27.1 ± 1.3<br>**Male *Mapt*$^{-/-}$ rats +YM+MTEP (10)**: 21.8 ± 3.7 | Two-way ANOVA:<br>**Interaction**: $F_{(1, 32)}$ = 9.1, *P* = 0.0051<br>**YM+MTEP effect**: $F_{(1, 32)}$ = 1.7, *P* = 0.20<br>**Genotype effect**: $F_{(1, 32)}$ = 1.2, *P* = 0.28 | Selected Šídák's post hoc test (adj. *P*-values):<br>**Control: WT vs. *Mapt*$^{-/-}$**: *P* = 0.026<br>**YM+MTEP: WT vs. *Mapt*$^{-/-}$**: *P* = 0.56<br>**WT: Control vs. YM+MTEP**: *P* = 0.67<br>***Mapt*$^{-/-}$: Control vs. YM+MTEP**: *P* = 0.018 |
| **Fig. 5e-g**. LTD+LatA/DMSO versus LTD+LatA/DMSO+YM+ MTEP in male WT rats | % LTD:<br>**Male WT rats +DMSO (6)**: 23.8 ± 2.0<br>**Male WT rats +LatA (8)**: 35.3 ± 2.5<br>**Male WT rats +DMSO+YM+MTEP (7)**: 20.6 ± 2.6<br>**Male WT rats +LatA+YM+MTEP (7)**: 19.9 ± 3.6 | Two-way ANOVA:<br>**Interaction**: $F_{(1, 24)}$ = 4.74 , *P* = 0.0394<br>**YM+MTEP effect**: $F_{(1, 24)}$ = 11.2, *P* = 0.0027<br>**LatA effect**: $F_{(1, 24)}$ = 3.80, *P* = 0.063 | Selected Šídák's post hoc test (adj. *P*-values):<br>**Control: DMSO vs. Lat A**: *P* = 0.031<br>**YM+MTEP: DMSO vs. Lat A**: *P* = 0.99<br>**DMSO: Control vs. YM+MTEP**: *P* = 0.9<br>**LatA: Control vs. YM+MTEP**: *P* = 0.0018 |
| **Fig. 5h-j**. LTD+LatA/DMSO versus LTD+LatA/DMSO+YM+ MTEP in male *Mapt*$^{-/-}$ rats | % LTD:<br>**Male *Mapt*$^{-/-}$ rats +DMSO (7)**: 32.6 ± 2.9<br>**Male *Mapt*$^{-/-}$ rats +LatA (7)**: 26.9 ± 2.9<br>**Male *Mapt*$^{-/-}$ rats +DMSO+YM+MTEP (7)**: 20.5 ± 2.0<br>**Male *Mapt*$^{-/-}$ rats +LatA+YM+MTEP (7)**: 23.0 ± 3.4 | Two-way ANOVA:<br>**Interaction**: $F_{(1, 24)}$ = 2.06 , *P* = 0.164<br>**YM+MTEP effect**: $F_{(1, 24)}$ = 7.76, *P* = 0.0102<br>**LatA effect**: $F_{(1, 24)}$ = 0.304, *P* = 0.586 | Selected Šídák's post hoc test (adj. *P*-values):<br>**Control: DMSO vs. Lat A**: *P* = 0.53<br>**YM+MTEP: DMSO vs. Lat A**: *P* = 0.95<br>**DMSO: Control vs. YM+MTEP**: *P* = 0.025<br>**LatA: Control vs. YM+MTEP**: *P* = 0.8 |
| **Extended Data Fig. 5m-o**. DHPG-LTD in male and female WT versus *Mapt*$^{-/-}$ rats | % LTD:<br>**Male WT rats (8)**: 21.3 ± 2.24<br>**Male *Mapt*$^{-/-}$ rats (8)**: 23.2 ± 2.15<br>**Female WT rats (6)**: 23.1 ± 3.23<br>**Female *Mapt*$^{-/-}$ rats (6)**: 22.4 ± 1.61 | Two-way ANOVA:<br>**Interaction**: $F_{(1, 24)}$ = 0.29, *P* = 0.58<br>**Sex effect**: $F_{(1, 24)}$ = 0.047, *P* = 0.83<br>**Genotype effect**: $F_{(1, 24)}$ = 0.071, *P* = 0.79 | N/A (no significant interactions, or main effects in two-way ANOVA) |
| **Extended Data Fig. 6a,b.** LTD+LatA/DMSO in female WT rats. | % LTD:<br>**Female WT rats +DMSO (7)**: 22.3 ± 4.2%<br>**Female WT rats +LatA (7)**: 25.8 ± 2.2% | Unpaired Student's *t*-test:<br>***t*-statistic**: 0.7<br>**Df**: 12<br>***P*-value**: 0.5 | N/A |
| **Extended Data Fig. 6c,d.** LTD+LatA/DMSO in female *Mapt*$^{-/-}$ rats. | % LTD:<br>**Female *Mapt*$^{-/-}$ rats +DMSO (6)**: 23.3 ± 2.6%<br>**Female *Mapt*$^{-/-}$ rats +LatA (6)**: 24.9 ± 3.9% | Unpaired Student's *t*-test:<br>***t*-statistic**: 0.3<br>**Df**: 10<br>***P*-value**: 0.7 | N/A |

**Supplementary Table 3 (part 1)**. Statistics summary for WT and *Mapt*$^{-/-}$ rat immunoblotting.

| Figure number and experiment: | Group (number of rodents) and mean ± SEM: | Statistical tests, *F/t*-statistic, degrees of freedoms (dfn,d), and *P*-values | Post-hoc test and *P*-values: |
|---|---|---|---|
| **Fig. 1b**. + **Extended Data Fig. 2a.** Total hippocampal homogenate tau levels | % expression (relative to male WT):<br>**Male WT rats (8)**: 100.0 ± 4.44<br>**Male *Mapt*$^{-/-}$ rats (8)**: 0.52 ± 0.10<br>**Female WT rats (8)**: 98.58 ± 3.37<br>**Female *Mapt*$^{-/-}$ rats (8)**: 0.32 ± 0.03 | Two-way ANOVA:<br>**Interaction**: $F_{(1, 28)}$ = 0.048, $P$ = 0.83<br>**Sex effect**: $F_{(1, 28)}$ = 0.084, $P$ = 0.78<br>**Genotype effect**: $F_{(1, 28)}$ = 1258 , $P$ < 0.0001 | Selected Šídák's post hoc test (adj. *P*-values):<br>**Male WT vs. Male *Mapt*$^{-/-}$**: $P$ < 0.0001<br>**Female WT vs. Female *Mapt*$^{-/-}$**: $P$ < 0.0001 |
| **Fig. 1b**. + **Extended Data Fig. 2b.** Total hippocampal homogenate MAP1A levels | % expression (relative to male WT):<br>**Male WT rats (8)**: 100.00 ± 11.04<br>**Male *Mapt*$^{-/-}$ rats (8)**: 94.29 ± 7.97<br>**Female WT rats (8)**: 92.79 ± 9.09<br>**Female *Mapt*$^{-/-}$ rats (8)**: 88.30 ± 12.15 | Two-way ANOVA:<br>**Interaction**: $F_{(1, 28)}$ = 0.0036, $P$ = 0.95<br>**Sex effect**: $F_{(1, 28)}$ = 0.42, $P$ = 0.52<br>**Genotype effect**: $F_{(1, 28)}$ = 0.25, $P$ = 0.62 | N/A (no significant interactions, or main effects in two-way ANOVA) |
| **Fig. 1b**. + **Extended Data Fig. 2c.** Total hippocampal homogenate MAP1B levels | % expression (relative to male WT):<br>**Male WT rats (8)**: 100.00 ± 13.76<br>**Male *Mapt*$^{-/-}$ rats (8)**: 97.78 ± 12.62<br>**Female WT rats (8)**: 93.11 ± 10.31<br>**Female *Mapt*$^{-/-}$ rats (8)**: 93.40 ± 12.64 | Two-way ANOVA:<br>**Interaction**: $F_{(1, 28)}$ = 0.01, $P$ = 0.92<br>**Sex effect**: $F_{(1, 28)}$ = 0.2, $P$ = 0.65<br>**Genotype effect**: $F_{(1, 28)}$ = 0.006, $P$ = 0.94 | N/A (no significant interactions, or main effects in two-way ANOVA) |
| **Fig. 1b**. + **Extended Data Fig. 2d.** Total hippocampal homogenate MAP2B levels | % expression (relative to male WT):<br>**Male WT rats (8)**: 100.00 ± 8.22<br>**Male *Mapt*$^{-/-}$ rats (8)**: 115.47 ± 10.86<br>**Female WT rats (8)**: 92.40 ± 11.80<br>**Female *Mapt*$^{-/-}$ rats (8)**: 105.36 ± 9.70 | Two-way ANOVA:<br>**Interaction**: $F_{(1, 28)}$ = 0.015, $P$ = 0.90<br>**Sex effect**: $F_{(1, 28)}$ = 0.75, $P$ = 0.39<br>**Genotype effect**: $F_{(1, 28)}$ = 1.93 , $P$ = 0.18 | N/A (no significant interactions, or main effects in two-way ANOVA) |
| **Fig. 1b**. + **Extended Data Fig. 2e.** Total hippocampal homogenate MAP2C levels | % expression (relative to male WT):<br>**Male WT rats (8)**: 100.00 ± 4.92<br>**Male *Mapt*$^{-/-}$ rats (8)**: 114.12 ± 6.00<br>**Female WT rats (8)**: 105.04 ± 4.43<br>**Female *Mapt*$^{-/-}$ rats (8)**: 108.84 ± 6.10 | Two-way ANOVA:<br>**Interaction**: $F_{(1, 28)}$ = 0.91, $P$ = 0.35<br>**Sex effect**: $F_{(1, 28)}$ = 0.00047, $P$ = 0.98<br>**Genotype effect**: $F_{(1, 28)}$ = 2.75, $P$ = 0.11 | N/A (no significant interactions, or main effects in two-way ANOVA) |
| **Fig. 1c**. + **Extended Data Fig. 2f.** Synaptically-enriched hippocampal tau levels | % expression (relative to male WT):<br>**Male WT rats (8)**: 100.00 ± 12.06<br>**Male *Mapt*$^{-/-}$ rats (8)**: 0.62 ± 0.15<br>**Female WT rats (8)**: 112.74 ± 10.93<br>**Female *Mapt*$^{-/-}$ rats (8)**: 0.81 ± 0.17 | Two-way ANOVA:<br>**Interaction**: $F_{(1, 28)}$ = 0.59, $P$ = 0.45<br>**Sex effect**: $F_{(1, 28)}$ = 0.63, $P$ = 0.43<br>**Genotype effect**: $F_{(1, 28)}$ = 168.5, $P$ < 0.0001 | Selected Šídák's post hoc test (adj. *P*-values):<br>**Male WT vs. Male *Mapt*$^{-/-}$**: $P$ < 0.0001<br>**Female WT vs. Female *Mapt*$^{-/-}$**: $P$ < 0.0001 |
| **Fig. 1c**. + **Extended Data Fig. 2g.** Synaptically-enriched hippocampal MAP1A levels | % expression (relative to male WT):<br>**Male WT rats (8)**: 100.00 ± 7.31<br>**Male *Mapt*$^{-/-}$ rats (8)**: 98.58 ± 9.94<br>**Female WT rats (8)**: 89.95 ± 6.64<br>**Female *Mapt*$^{-/-}$ rats (8)**: 85.91 ± 7.05 | Two-way ANOVA:<br>**Interaction**: $F_{(1, 28)}$ = 0.028, $P$ = 0.87<br>**Sex effect**: $F_{(1, 28)}$ = 2.1, $P$ = 0.16<br>**Genotype effect**: $F_{(1, 28)}$ = 0.12, $P$ = 0.73 | N/A (no significant interactions, or main effects in two-way ANOVA) |
| **Fig. 1c**. + **Extended Data Fig. 2h.** Synaptically-enriched hippocampal MAP1B levels | % expression (relative to male WT):<br>**Male WT rats (8)**: 100.00 ± 13.94<br>**Male *Mapt*$^{-/-}$ rats (8)**: 88.44 ± 15.49<br>**Female WT rats (8)**: 82.05 ± 9.98<br>**Female *Mapt*$^{-/-}$ rats (8)**: 84.77 ± 12.14 | Two-way ANOVA:<br>**Interaction**: $F_{(1, 28)}$ = 0.3, $P$ = 0.59<br>**Sex effect**: $F_{(1, 28)}$ = 0.69, $P$ = 0.42<br>**Genotype effect**: $F_{(1, 28)}$ = 0.12 , $P$ = 0.74 | N/A (no significant interactions, or main effects in two-way ANOVA) |
| **Fig. 1c**. + **Extended Data Fig. 2i.** Synaptically-enriched hippocampal MAP2B levels | % expression (relative to male WT):<br>**Male WT rats (8)**: 100.00 ± 8.46<br>**Male *Mapt*$^{-/-}$ rats (8)**: 123.32 ± 17.99<br>**Female WT rats (8)**: 99.54 ± 9.56<br>**Female *Mapt*$^{-/-}$ rats (8)**: 115.40 ± 13.45 | Two-way ANOVA:<br>**Interaction**: $F_{(1, 28)}$ = 0.083, $P$ = 0.78<br>**Sex effect**: $F_{(1, 28)}$ = 0.11, $P$ = 0.75<br>**Genotype effect**: $F_{(1, 28)}$ = 2.3, $P$ = 0.14 | N/A (no significant interactions, or main effects in two-way ANOVA) |
| **Fig. 1c**. + **Extended Data Fig. 2j.** Synaptically-enriched hippocampal MAP2C levels | % expression (relative to male WT):<br>**Male WT rats (8)**: 100.00 ± 12.88<br>**Male *Mapt*$^{-/-}$ rats (8)**: 104.95 ± 12.99<br>**Female WT rats (8)**: 98.54 ± 11.09<br>**Female *Mapt*$^{-/-}$ rats (8)**: 105.68 ± 14.93 | Two-way ANOVA:<br>**Interaction**: $F_{(1, 28)}$ = 0.007, $P$ = 0.93<br>**Sex effect**: $F_{(1, 28)}$ = 0.00077, $P$ = 0.98<br>**Genotype effect**: $F_{(1, 28)}$ = 0.22, $P$ = 0.65 | N/A (no significant interactions, or main effects in two-way ANOVA) |
| **Extended Data Fig. 4a.** Synaptically-enriched hippocampal GluN2A levels | % expression (relative to male WT):<br>**Male WT rats (8)**: 100.00 ± 14.41<br>**Male *Mapt*$^{-/-}$ rats (8)**: 94.57 ± 11.15<br>**Female WT rats (8)**: 154.11 ± 20.01<br>**Female *Mapt*$^{-/-}$ rats (8)**: 129.56 ± 17.54 | Two-way ANOVA:<br>**Interaction**: $F_{(1, 28)}$ = 0.35, $P$ = 0.56<br>**Sex effect**: $F_{(1, 28)}$ = 7.63, $P$ = 0.01<br>**Genotype effect**: $F_{(1, 28)}$ = 0.86, $P$ = 0.36 | Selected Šídák's post hoc test (adj. *P*-values):<br>**Male WT vs. Female WT**: $P$ = 0.023 |
| **Extended Data Fig. 4b.** Synaptically-enriched hippocampal pGluN2A Y1426 levels | % expression (relative to male WT):<br>**Male WT rats (8)**: 100.00 ± 15.31<br>**Male *Mapt*$^{-/-}$ rats (8)**: 109.39 ± 19.50<br>**Female WT rats (8)**: 74.39 ± 15.69<br>**Female *Mapt*$^{-/-}$ rats (8)**: 97.09 ± 18.35 | Two-way ANOVA:<br>**Interaction**: $F_{(1, 28)}$ = 0.15, $P$ = 0.70<br>**Sex effect**: $F_{(1, 28)}$ = 1.2, $P$ = 0.28<br>**Genotype effect**: $F_{(1, 28)}$ = 0.86, $P$ = 0.36 | N/A (no significant interactions, or main effects in two-way ANOVA) |
| **Extended Data Fig. 4c.** Synaptically-enriched hippocampal GluN2B levels | % expression (relative to male WT):<br>**Male WT rats (8)**: 100.00 ± 7.87<br>**Male *Mapt*$^{-/-}$ rats (8)**: 98.20 ± 6.65<br>**Female WT rats (8)**: 94.14 ± 8.81<br>**Female *Mapt*$^{-/-}$ rats (8)**: 109.12 ± 4.36 | Two-way ANOVA:<br>**Interaction**: $F_{(1, 28)}$ = 1.4, $P$ = 0.25<br>**Sex effect**: $F_{(1, 28)}$ = 0.13, $P$ = 0.73<br>**Genotype effect**: $F_{(1, 28)}$ = 0.86, $P$ = 0.36 | N/A (no significant interactions, or main effects in two-way ANOVA) |
| **Extended Data Fig. 4d.** Synaptically-enriched hippocampal pGluN2B Y1472 levels | % expression (relative to male WT):<br>**Male WT rats (8)**: 100.00 ± 12.38<br>**Male *Mapt*$^{-/-}$ rats (8)**: 79.51 ± 10.05<br>**Female WT rats (8)**: 124.21 ± 11.47<br>**Female *Mapt*$^{-/-}$ rats (8)**: 115.51 ± 13.33 | Two-way ANOVA:<br>**Interaction**: $F_{(1, 28)}$ = 0.25, $P$ = 0.62<br>**Sex effect**: $F_{(1, 28)}$ = 6.43, $P$ = 0.02<br>**Genotype effect**: $F_{(1, 28)}$ = 1.51, $P$ = 0.23 | Selected Šídák's post hoc test (adj. *P*-values):<br>**Male WT vs. Female WT**: $P$ = 0.3 |
| **Extended Data Fig. 4e.** Synaptically-enriched hippocampal GluA1 levels | % expression (relative to male WT):<br>**Male WT rats (8)**: 100.00 ± 6.25<br>**Male *Mapt*$^{-/-}$ rats (6)**: 99.30 ± 10.02<br>**Female WT rats (8)**: 101.93 ± 7.15<br>**Female *Mapt*$^{-/-}$ rats (5)**: 95.94 ± 13.98 | Two-way ANOVA:<br>**Interaction**: $F_{(1, 28)}$ = 0.087, $P$ = 0.77<br>**Sex effect**: $F_{(1, 28)}$ = 0.0065, $P$ = 0.94<br>**Genotype effect**: $F_{(1, 28)}$ = 0.14, $P$ = 0.71 | N/A (no significant interactions, or main effects in two-way ANOVA) |

**Supplementary Table 3 (part 2)**. Statistics summary for WT and *Mapt*$^{-/-}$ rat immunoblotting.

| **Figure number and experiment:** | **Group (number of rodents) and mean ± SEM:** | **Statistical tests, *F/t*-statistic, degrees of freedoms (dfn,d), and *P*-values** | **Post-hoc test and *P*-values:** |
|---|---|---|---|
| **Extended Data Fig. 4f.** Synaptically-enriched hippocampal pGluA1 S845 levels | % expression (relative to male WT):<br>**Male WT rats (8)**: 100.00 ± 13.81<br>**Male *Mapt*$^{-/-}$ rats (6)**: 120.03 ± 11.12<br>**Female WT rats (8)**: 114.14 ± 8.01<br>**Female *Mapt*$^{-/-}$ rats (4)**: 115.65 ± 27.68 | Two-way ANOVA:<br>**Interaction**: $F_{(1, 28)}$ =0.42, $P$ = 0.53<br>**Sex effect**: $F_{(1, 28)}$ = 0.12, $P$ = 0.74<br>**Genotype effect**: $F_{(1, 28)}$ = 0.56, $P$ = 0.46 | N/A (no significant interactions, or main effects in two-way ANOVA) |
| **Extended Data Fig. 4g.** Synaptically-enriched hippocampal GluA2 levels | % expression (relative to male WT):<br>**Male WT rats (8)**: 100.00 ± 10.83<br>**Male *Mapt*$^{-/-}$ rats (7)**: 120.11 ± 15.48<br>**Female WT rats (7)**: 117.90 ± 12.68<br>**Female *Mapt*$^{-/-}$ rats (8)**: 130.94 ± 16.36 | Two-way ANOVA:<br>**Interaction**: $F_{(1, 28)}$ = 0.063, $P$ = 0.80<br>**Sex effect**: $F_{(1, 28)}$ = 1.04, $P$ = 0.32<br>**Genotype effect**: $F_{(1, 28)}$ = 0.139, $P$ = 0.25 | N/A (no significant interactions, or main effects in two-way ANOVA) |
| **Extended Data Fig. 4h.** Synaptically-enriched hippocampal pGluA2 S880 levels | % expression (relative to male WT):<br>**Male WT rats (7)**: 100.00 ± 13.58<br>**Male *Mapt*$^{-/-}$ rats (6)**: 94.83 ± 12.25<br>**Female WT rats (5)**: 79.79 ± 5.62<br>**Female *Mapt*$^{-/-}$ rats (8)**: 96.31 ± 11.66 | Two-way ANOVA:<br>**Interaction**: $F_{(1, 28)}$ = 0.80, $P$ = 0.38<br>**Sex effect**: $F_{(1, 28)}$ = 0.59, $P$ = 0.45<br>**Genotype effect**: $F_{(1, 28)}$ = 0.22, $P$ = 0.65 | N/A (no significant interactions, or main effects in two-way ANOVA) |
| **Extended Data Fig. 5a,b.** Total hippocampal mGlu1 dimer levels | % expression (relative to male WT):<br>**Male WT rats (8)**: 100.00 ± 14.64<br>**Male *Mapt*$^{-/-}$ rats (8)**: 104.69 ± 13.82<br>**Female WT rats (8)**: 99.77 ± 11.73<br>**Female *Mapt*$^{-/-}$ rats (8)**: 98.15 ± 11.39 | Two-way ANOVA:<br>**Interaction**: $F_{(1, 28)}$ = 0.059, $P$ = 0.81<br>**Sex effect**: $F_{(1, 28)}$ = 0.068, $P$ = 0.8<br>**Genotype effect**: $F_{(1, 28)}$ = 0.014, $P$ = 0.91 | N/A (no significant interactions, or main effects in two-way ANOVA) |
| **Extended Data Fig. 5a,c.** Total hippocampal mGlu1 monomer levels | % expression (relative to male WT):<br>**Male WT rats (8)**: 100.00 ± 13.60<br>**Male *Mapt*$^{-/-}$ rats (8)**: 107.35 ± 11.37<br>**Female WT rats (8)**: 104.62 ± 9.04<br>**Female *Mapt*$^{-/-}$ rats (8)**: 95.04 ± 10.09 | Two-way ANOVA:<br>**Interaction**: $F_{(1, 28)}$ = 0.58, $P$ = 0.45<br>**Sex effect**: $F_{(1, 28)}$ = 0.12, $P$ = 0.73<br>**Genotype effect**: $F_{(1, 28)}$ = 0.001, $P$ = 0.92 | N/A (no significant interactions, or main effects in two-way ANOVA) |
| **Extended Data Fig. 5d,e.** Total hippocampal mGlu5 dimer levels | % expression (relative to male WT):<br>**Male WT rats (8)**: 100.00 ± 14.33<br>**Male *Mapt*$^{-/-}$ rats (8)**: 105.55 ± 11.70<br>**Female WT rats (8)**: 92.49 ± 11.25<br>**Female *Mapt*$^{-/-}$ rats (8)**: 83.16 ± 8.31 | Two-way ANOVA:<br>**Interaction**: $F_{(1, 28)}$ = 0.41, $P$ = 0.53<br>**Sex effect**: $F_{(1, 28)}$ = 1.66, $P$ = 0.21<br>**Genotype effect**: $F_{(1, 28)}$ = 0.027, $P$ = 0.87 | N/A (no significant interactions, or main effects in two-way ANOVA) |
| **Extended Data Fig. 5d,f.** Total hippocampal mGlu5 monomer levels | % expression (relative to male WT):<br>**Male WT rats (8)**: 100.00 ± 9.40<br>**Male *Mapt*$^{-/-}$ rats (8)**: 110.93 ± 9.19<br>**Female WT rats (8)**: 100.91 ± 8.34<br>**Female *Mapt*$^{-/-}$ rats (8)**: 99.61 ± 7.50 | Two-way ANOVA:<br>**Interaction**: $F_{(1, 28)}$ = 0.5, $P$ = 0.49<br>**Sex effect**: $F_{(1, 28)}$ = 0.36, $P$ = 0.55<br>**Genotype effect**: $F_{(1, 28)}$ = 0.31 $P$ = 0.58 | N/A (no significant interactions, or main effects in two-way ANOVA) |
| **Extended Data Fig. 5g,h.** Synaptically-enriched hippocampal mGlu1 dimer levels | % expression (relative to male WT):<br>**Male WT rats (8)**: 100.00 ± 13.46<br>**Male *Mapt*$^{-/-}$ rats (8)**: 91.30 ± 9.05<br>**Female WT rats (8)**: 102.50 ± 7.60<br>**Female *Mapt*$^{-/-}$ rats (8)**: 104.02 ± 6.85 | Two-way ANOVA:<br>**Interaction**: $F_{(1, 28)}$ = 0.28, $P$ = 0.6<br>**Sex effect**: $F_{(1, 28)}$ = 0.63, $P$ = 0.43<br>**Genotype effect**: $F_{(1, 28)}$ = 0.14, $P$ = 0.71 | N/A (no significant interactions, or main effects in two-way ANOVA) |
| **Extended Data Fig. 5g,i.** Synaptically-enriched hippocampal mGlu1 monomer levels | % expression (relative to male WT):<br>**Male WT rats (8)**: 100.00 ± 13.79<br>**Male *Mapt*$^{-/-}$ rats (8)**: 91.28 ± 15.04<br>**Female WT rats (8)**: 106.06 ± 6.74<br>**Female *Mapt*$^{-/-}$ rats (8)**: 109.02 ± 11.46 | Two-way ANOVA:<br>**Interaction**: $F_{(1, 28)}$ = 0.23, $P$ = 0.64<br>**Sex effect**: $F_{(1, 28)}$ = 0.96, $P$ = 0.34<br>**Genotype effect**: $F_{(1, 28)}$ = 0.056, $P$ = 0.82 | N/A (no significant interactions, or main effects in two-way ANOVA) |
| **Extended Data Fig. 5j,k.** Synaptically-enriched hippocampal mGlu5 dimer levels | % expression (relative to male WT):<br>**Male WT rats (8)**: 100.00 ± 7.03<br>**Male *Mapt*$^{-/-}$ rats (8)**: 105.51 ± 5.71<br>**Female WT rats (8)**: 108.62 ± 9.02<br>**Female *Mapt*$^{-/-}$ rats (8)**: 104.37 ± 8.55 | Two-way ANOVA:<br>**Interaction**: $F_{(1, 28)}$ = 0.4, $P$ = 0.53<br>**Sex effect**: $F_{(1, 28)}$ = 0.24, $P$ = 0.63<br>**Genotype effect**: $F_{(1, 28)}$ = 0.0067, $P$ = 0.94 | N/A (no significant interactions, or main effects in two-way ANOVA) |
| **Extended Data Fig. 5j,l.** Synaptically-enriched hippocampal mGlu5 monomer levels | % expression (relative to male WT):<br>**Male WT rats (8)**: 100.00 ± 14.58<br>**Male *Mapt*$^{-/-}$ rats (8)**: 94.54 ± 12.81<br>**Female WT rats (8)**: 90.39 ± 10.15<br>**Female *Mapt*$^{-/-}$ rats (8)**: 87.83 ± 8.98 | Two-way ANOVA:<br>**Interaction**: $F_{(1, 28)}$ = 0.015, $P$ = 0.9<br>**Sex effect**: $F_{(1, 28)}$ = 0.48, $P$ = 0.5<br>**Genotype effect**: $F_{(1, 28)}$ = 0.12, $P$ = 0.74 | N/A (no significant interactions, or main effects in two-way ANOVA) |
| **Fig.5a** Total hippocampal homogenate α-tubulin levels | % expression (relative to male WT):<br>**Male WT rats (8)**: 100.00 ± 6.09<br>**Male *Mapt*$^{-/-}$ rats (8)**: 102.14 ± 3.27<br>**Female WT rats (8)**: 104.23 ± 7.05<br>**Female *Mapt*$^{-/-}$ rats (8)**: 99.13 ± 6.08 | Two-way ANOVA:<br>**Interaction**: $F_{(1, 28)}$ = 0.39, $P$ = 0.54<br>**Sex effect**: $F_{(1, 28)}$ = 0.011, $P$ = 0.92<br>**Genotype effect**: $F_{(1, 28)}$ = 0.065, $P$ = 0.8 | N/A (no significant interactions, or main effects in two-way ANOVA) |
| **Fig. 5b** Total hippocampal homogenate β-actin levels | % expression (relative to male WT):<br>**Male WT rats (8)**: 100.00 ± 4.33<br>**Male *Mapt*$^{-/-}$ rats (8)**: 103.21± 7.39<br>**Female WT rats (8)**: 96.95 ± 7.30<br>**Female *Mapt*$^{-/-}$ rats (8)**: 103.38 ± 10.64 | Two-way ANOVA:<br>**Interaction**: $F_{(1, 28)}$ = 0.043, $P$ = 0.84<br>**Sex effect**: $F_{(1, 28)}$ = 0.035, $P$ = 0.85<br>**Genotype effect**: $F_{(1, 28)}$ = 0.39, $P$ = 0.54 | N/A (no significant interactions, or main effects in two-way ANOVA) |
| **Fig. 5c** Synaptically-enriched hippocampal α-tubulin levels | % expression (relative to male WT):<br>**Male WT rats (8)**: 100.00 ± 8.59<br>**Male *Mapt*$^{-/-}$ rats (8)**: 108.13 ± 16.54<br>**Female WT rats (8)**: 98.31 ± 12.52<br>**Female *Mapt*$^{-/-}$ rats (8)**: 91.29 ± 8.79 | Two-way ANOVA:<br>**Interaction**: $F_{(1, 28)}$ = 0.4, $P$ = 0.54<br>**Sex effect**: $F_{(1, 28)}$ = 0.59, $P$ = 0.45<br>**Genotype effect**: $F_{(1, 28)}$ = 0.0021, $P$ = 0.96 | N/A (no significant interactions, or main effects in two-way ANOVA) |
| **Fig. 5d** Synaptically-enriched hippocampal β-actin levels | % expression (relative to male WT):<br>**Male WT rats (8)**: 100.00 ± 5.94<br>**Male *Mapt*$^{-/-}$ rats (8)**: 131.87 ± 12.83<br>**Female WT rats (8)**: 100.92 ± 6.01<br>**Female *Mapt*$^{-/-}$ rats (8)**: 137.18 ± 8.45 | Two-way ANOVA:<br>**Interaction**: $F_{(1, 28)}$ = 0.0066, $P$ = 0.94<br>**Sex effect**: $F_{(1, 28)}$ = 0.083, $P$ = 0.78<br>**Genotype effect**: $F_{(1, 28)}$ = 11.66, $P$ = 0.002 | Selected Šídák's post hoc test (adj. *P*-values):<br>**Male WT vs. *Mapt*$^{-/-}$**: $P$ = 0.031<br>**Female WT vs. *Mapt*$^{-/-}$**: $P$ = 0.014 |

**Supplementary Figure 1**. Lane order, full-blot images, total protein images, and primary antibody targets for representative immunoblots for *Mapt*$^{-/-}$ rat immunoblotting experiments.

**a**

### Total tau (hippocampal homogenates)

| Set 3 of 4 | |
|---|---|
| **Lane #:** | **Sample** |
| 1 | Size marker |
| 2 | ♂ *Mapt*−/− ○ |
| 3 | ♂ WT ● |
| 4 | ♂ WT ● |
| 5 | ♂ *Mapt*−/− ○ |
| 6 | ♀ WT ▲ |
| 7 | ♀ *Mapt*−/− △ |
| 8 | ♀ *Mapt*−/− △ |
| 9 | ♀ WT ▲ |
| 10 | Internal control |

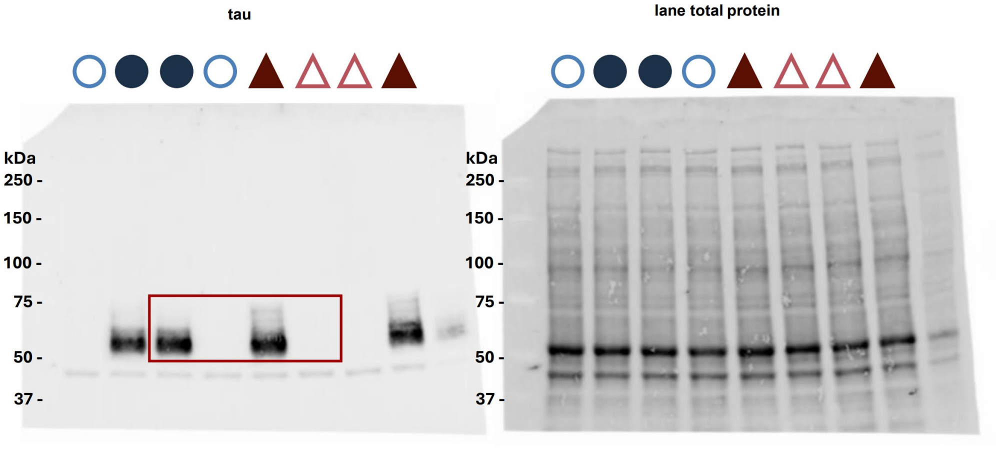


for Fig. 1b/Extended Data Fig. 2a

**b**

### Total MAP1A/ MAP1B (hippocampal homogenates)

| Set 1 of 4 | |
|---|---|
| **Lane #:** | **Sample** |
| 1 | Size marker |
| 2 | ♂ *Mapt*−/− ○ |
| 3 | ♂ WT ● |
| 4 | ♂ WT ● |
| 5 | ♂ *Mapt*−/− ○ |
| 6 | ♀ WT ▲ |
| 7 | ♀ *Mapt*−/− △ |
| 8 | ♀ WT ▲ |
| 9 | ♀ *Mapt*−/− △ |
| 10 | Internal control |

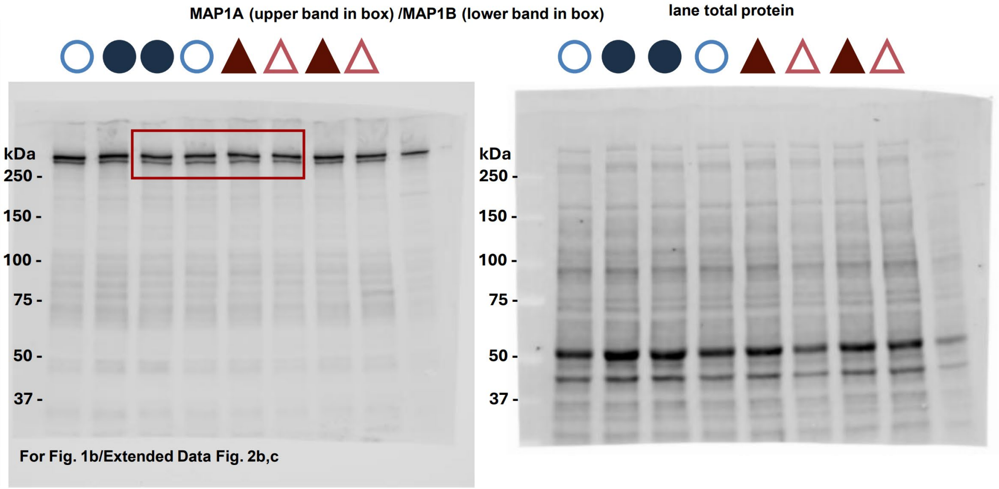


For Fig. 1b/Extended Data Fig. 2b,c

**c**

### Total MAP2B/ MAP2C (hippocampal homogenates)

| Set 1 of 4 | |
|---|---|
| **Lane #:** | **Sample** |
| 1 | Size marker |
| 2 | ♂ *Mapt*−/− ○ |
| 3 | ♂ WT ● |
| 4 | ♂ WT ● |
| 5 | ♂ *Mapt*−/− ○ |
| 6 | ♀ WT ▲ |
| 7 | ♀ *Mapt*−/− △ |
| 8 | ♀ WT ▲ |
| 9 | ♀ *Mapt*−/− △ |
| 10 | Internal control |

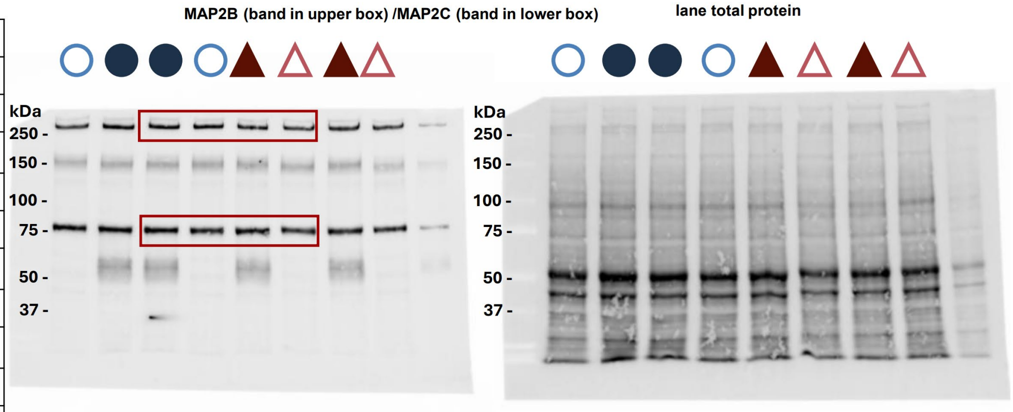


for Fig. 1b/Extended Data Fig. 2d,e

**d**

### Synaptic tau (hippocampal homogenates)

| Set 3 of 4 | |
|---|---|
| Lane #: | Sample |
| 1 | Size marker |
| 2 | ♂ *Mapt*$^{-/-}$ |
| 3 | ♂ WT |
| 4 | ♂ WT |
| 5 | ♂ *Mapt*$^{-/-}$ |
| 6 | ♀ WT |
| 7 | ♀ *Mapt*$^{-/-}$ |
| 8 | ♀ *Mapt*$^{-/-}$ |
| 9 | ♀ WT |
| 10 | Internal control |

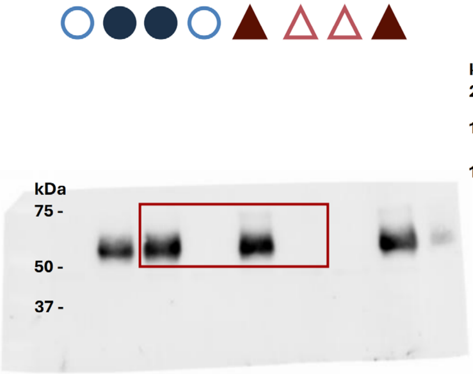


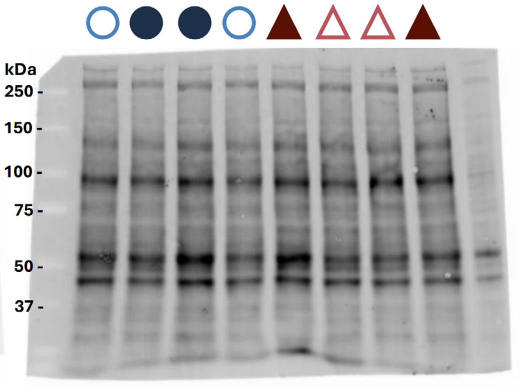


for Fig. 1c/Extended Data Fig. 2f

**e**

### Synaptic MAP1A/ MAP1B (hippocampal homogenates)

| Set 1 of 4 | |
|---|---|
| Lane #: | Sample |
| 1 | Size marker |
| 2 | ♂ *Mapt*$^{-/-}$ |
| 3 | ♂ WT |
| 4 | ♂ WT |
| 5 | ♂ *Mapt*$^{-/-}$ |
| 6 | ♀ WT |
| 7 | ♀ *Mapt*$^{-/-}$ |
| 8 | ♀ WT |
| 9 | ♀ *Mapt*$^{-/-}$ |
| 10 | Internal control |

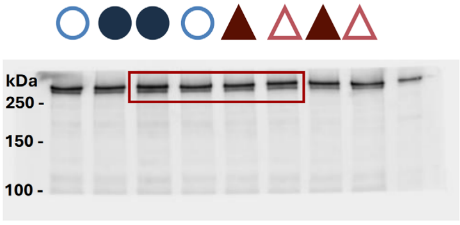


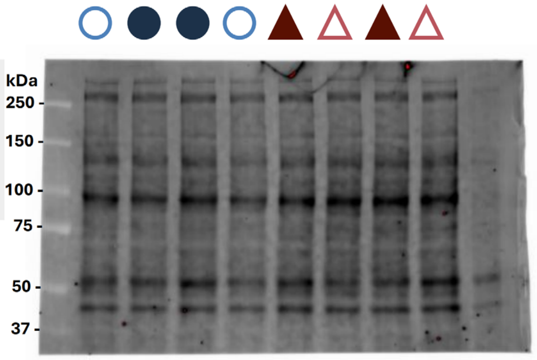


for Fig. 1c/Extended Data Fig. 2g,h

**f**

### Synaptic MAP2B/ MAP2C (hippocampal homogenates)

| Set 3 of 4 | |
|---|---|
| Lane #: | Sample |
| 1 | Size marker |
| 2 | ♂ *Mapt*$^{-/-}$ |
| 3 | ♂ WT |
| 4 | ♂ WT |
| 5 | ♂ *Mapt*$^{-/-}$ |
| 6 | ♀ WT |
| 7 | ♀ *Mapt*$^{-/-}$ |
| 8 | ♀ *Mapt*$^{-/-}$ |
| 9 | ♀ WT |
| 10 | Internal control |

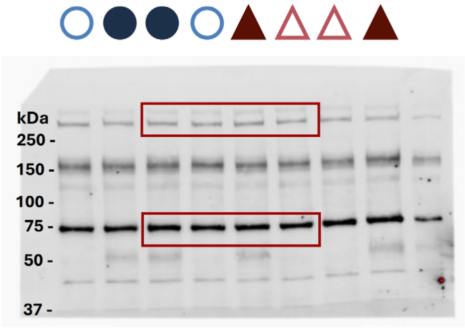


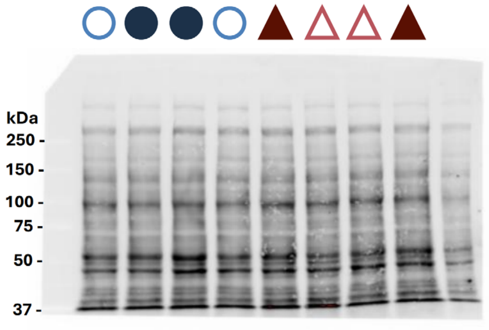


For Fig. 1c/Extended Data Fig. 2i,j

## g

### Synaptic GluN2A (hippocampal homogenates)

| Set 1 of 4 | |
|---|---|
| Lane #: | Sample |
| 1 | Size marker |
| 2 | ♂ *Mapt*$^{-/-}$ |
| 3 | ♂ WT |
| 4 | ♂ WT |
| 5 | ♂ *Mapt*$^{-/-}$ |
| 6 | ♀ WT |
| 7 | ♀ *Mapt*$^{-/-}$ |
| 8 | ♀ WT |
| 9 | ♀ *Mapt*$^{-/-}$ |
| 10 | Internal control |

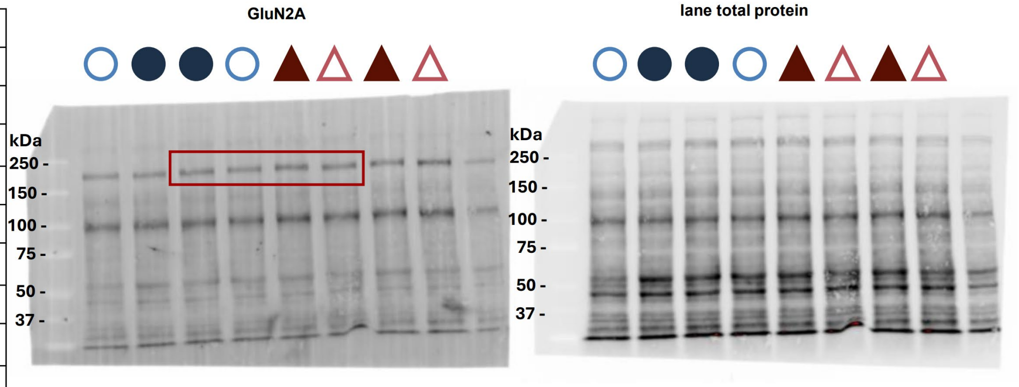


for Extended Data Fig. 4a

## h

### Synaptic pGluN2A Y1426 (hippocampal homogenates)

| Set 1 of 4 | |
|---|---|
| Lane #: | Sample |
| 1 | Size marker |
| 2 | ♂ *Mapt*$^{-/-}$ |
| 3 | ♂ WT |
| 4 | ♂ WT |
| 5 | ♂ *Mapt*$^{-/-}$ |
| 6 | ♀ WT |
| 7 | ♀ *Mapt*$^{-/-}$ |
| 8 | ♀ WT |
| 9 | ♀ *Mapt*$^{-/-}$ |
| 10 | Internal control |

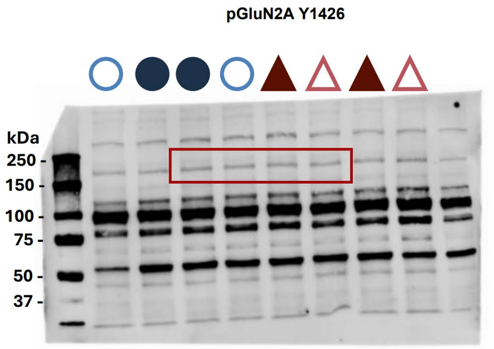


lane total protein

(see Synaptic GluN2A in panel g)

for Extended Data Fig. 4b

## i

### Synaptic GluN2B (hippocampal homogenates)

| Set 3 of 4 | |
|---|---|
| Lane #: | Sample |
| 1 | Size marker |
| 2 | ♂ *Mapt*$^{-/-}$ |
| 3 | ♂ WT |
| 4 | ♂ WT |
| 5 | ♂ *Mapt*$^{-/-}$ |
| 6 | ♀ WT |
| 7 | ♀ *Mapt*$^{-/-}$ |
| 8 | ♀ *Mapt*$^{-/-}$ |
| 9 | ♀ WT |
| 10 | Internal control |

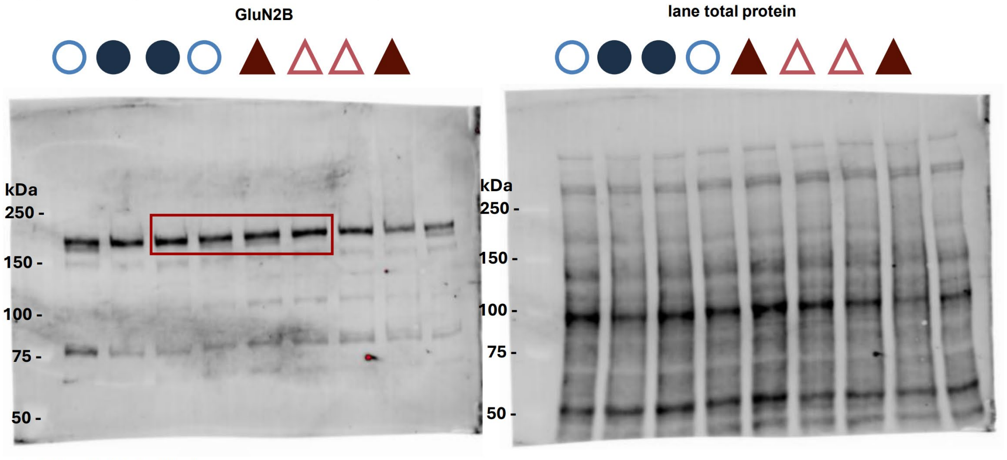


for Extended Data Fig. 4c

**j**

### Synaptic pGluN2B Y1472 (hippocampal homogenates)

| Set 3 of 4 | |
|---|---|
| **Lane #:** | **Sample** |
| 1 | Size marker |
| 2 | ♂ *Mapt*$^{-/-}$ ○ |
| 3 | ♂ WT ● |
| 4 | ♂ WT ● |
| 5 | ♂ *Mapt*$^{-/-}$ ○ |
| 6 | ♀ WT ▲ |
| 7 | ♀ *Mapt*$^{-/-}$ △ |
| 8 | ♀ *Mapt*$^{-/-}$ △ |
| 9 | ♀ WT ▲ |
| 10 | Internal control |

pGluN2B Y1472

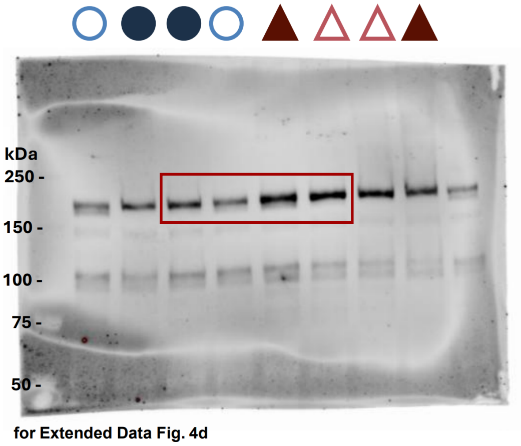


for Extended Data Fig. 4d

lane total protein

(see Synaptic GluN2B in panel i)

**k**

### Synaptic GluA1 (hippocampal homogenates)

| Set 1 of 4 | |
|---|---|
| **Lane #:** | **Sample** |
| 1 | Size marker |
| 2 | ♂ *Mapt*$^{-/-}$ ○ |
| 3 | ♂ WT ● |
| 4 | ♂ WT ● |
| 5 | ♂ *Mapt*$^{-/-}$ ○ |
| 6 | ♀ WT ▲ |
| 7 | ♀ *Mapt*$^{-/-}$ △ |
| 8 | ♀ WT ▲ |
| 9 | ♀ *Mapt*$^{-/-}$ △ |
| 10 | Internal control |

GluA1

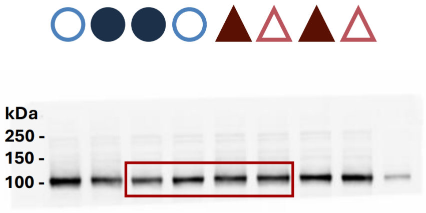


for Extended Data Fig. 4e

lane total protein

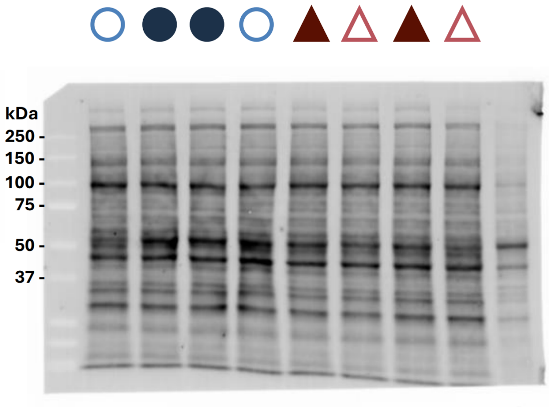


**l**

### Synaptic pGluA1 S845 (hippocampal homogenates)

| Set 1 of 4 | |
|---|---|
| **Lane #:** | **Sample** |
| 1 | Size marker |
| 2 | ♂ *Mapt*$^{-/-}$ ○ |
| 3 | ♂ WT ● |
| 4 | ♂ WT ● |
| 5 | ♂ *Mapt*$^{-/-}$ ○ |
| 6 | ♀ WT ▲ |
| 7 | ♀ *Mapt*$^{-/-}$ △ |
| 8 | ♀ WT ▲ |
| 9 | ♀ *Mapt*$^{-/-}$ △ |
| 10 | Internal control |

pGluA1 S845

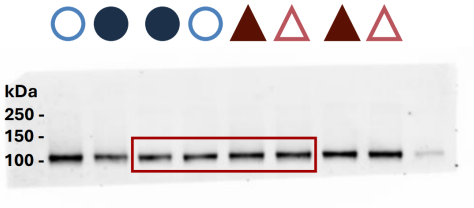


for Extended Data Fig. 4f

lane total protein

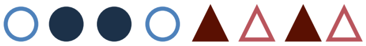

(see Synaptic GluA1 in panel k)

**m**

### Synaptic GluA2 (hippocampal homogenates)

| Set 3 of 4 | |
|---|---|
| Lane #: | Sample |
| 1 | Size marker |
| 2 | ♂ *Mapt*$^{-/-}$ |
| 3 | ♂ WT |
| 4 | ♂ WT |
| 5 | ♂ *Mapt*$^{-/-}$ |
| 6 | ♀ WT |
| 7 | ♀ *Mapt*$^{-/-}$ |
| 8 | ♀ *Mapt*$^{-/-}$ |
| 9 | ♀ WT |
| 10 | Internal control |

GluA2

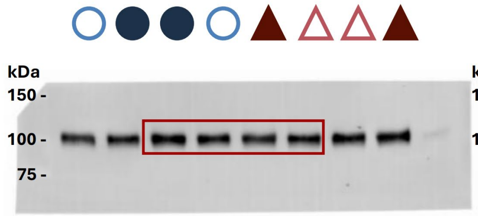


lane total protein

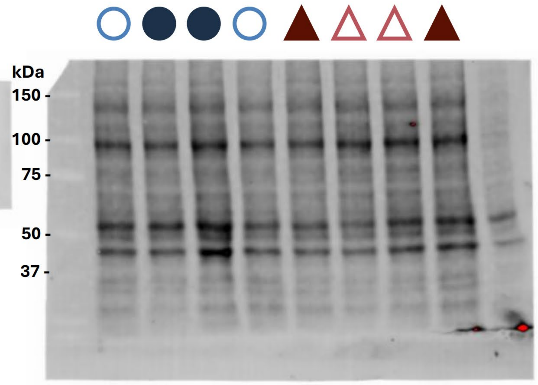


for Extended Data Fig. 4g

**n**

### Synaptic pGluA2 S880 (hippocampal homogenates)

| Set 3 of 4 | |
|---|---|
| Lane #: | Sample |
| 1 | Size marker |
| 2 | ♂ *Mapt*$^{-/-}$ |
| 3 | ♂ WT |
| 4 | ♂ WT |
| 5 | ♂ *Mapt*$^{-/-}$ |
| 6 | ♀ WT |
| 7 | ♀ *Mapt*$^{-/-}$ |
| 8 | ♀ *Mapt*$^{-/-}$ |
| 9 | ♀ WT |
| 10 | Internal control |

pGluA2 S880

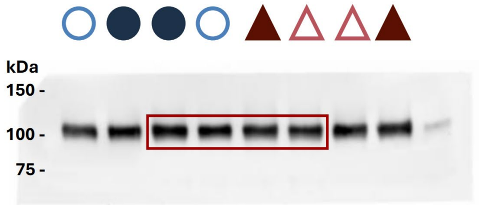


lane total protein

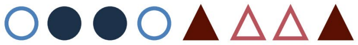

(see Synaptic GluA2 in panel m)

for Extended Data Fig. 4h

**o**

### Total mGlu1 monomers and dimers (hippocampal homogenates)

| Set 3 of 4 | |
|---|---|
| Lane #: | Sample |
| 1 | Size marker |
| 2 | ♂ *Mapt*$^{-/-}$ |
| 3 | ♂ WT |
| 4 | ♂ WT |
| 5 | ♂ *Mapt*$^{-/-}$ |
| 6 | ♀ WT |
| 7 | ♀ *Mapt*$^{-/-}$ |
| 8 | ♀ *Mapt*$^{-/-}$ |
| 9 | ♀ WT |
| 10 | Internal control |

mGlu1

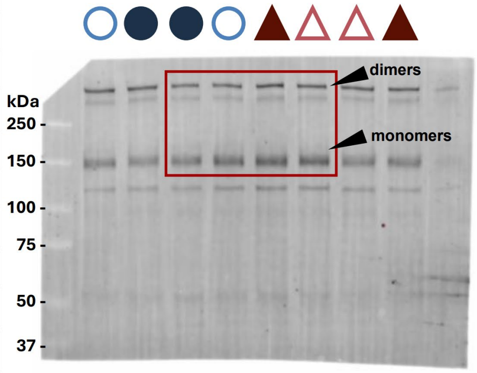


lane total protein

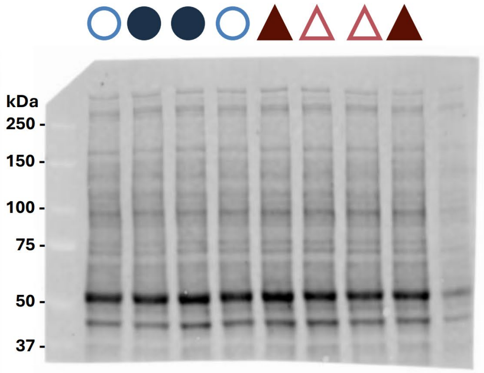


for Extended Fig. 5a-c

**p**

### Total mGlu5 monomers and dimers (hippocampal homogenates)

| Set 3 of 4 | |
|---|---|
| Lane #: | Sample |
| 1 | Size marker |
| 2 | ♂ *Mapt*$^{-/-}$ ○ |
| 3 | ♂ WT ● |
| 4 | ♂ WT ● |
| 5 | ♂ *Mapt*$^{-/-}$ ○ |
| 6 | ♀ WT ▲ |
| 7 | ♀ *Mapt*$^{-/-}$ △ |
| 8 | ♀ *Mapt*$^{-/-}$ △ |
| 9 | ♀ WT ▲ |
| 10 | Internal control |

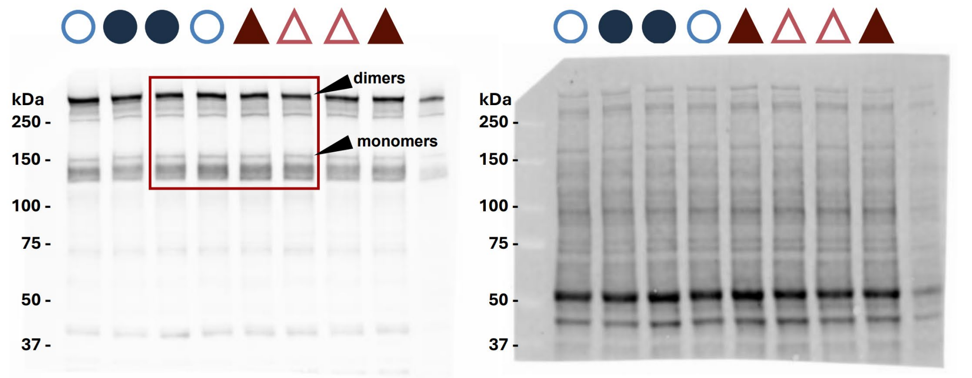


(same membrane as in panel o due to multiplexing antibodies)

for Extended Fig. 5d-f

**q**

### Synaptic mGlu1 monomers and dimers (hippocampal homogenates)

| Set 3 of 4 | |
|---|---|
| Lane #: | Sample |
| 1 | Size marker |
| 2 | ♂ *Mapt*$^{-/-}$ ○ |
| 3 | ♂ WT ● |
| 4 | ♂ WT ● |
| 5 | ♂ *Mapt*$^{-/-}$ ○ |
| 6 | ♀ WT ▲ |
| 7 | ♀ *Mapt*$^{-/-}$ △ |
| 8 | ♀ *Mapt*$^{-/-}$ △ |
| 9 | ♀ WT ▲ |
| 10 | Internal control |

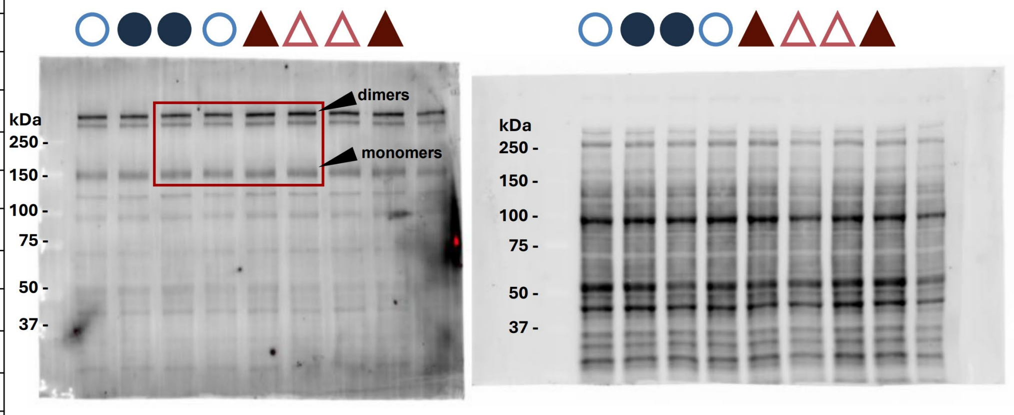


for Extended Fig. 5g-i

**r**

### Synaptic mGlu5 monomers + dimers (hippocampal homogenates)

| Set 1 of 4 | |
|---|---|
| Lane #: | Sample |
| 1 | Size marker |
| 2 | ♂ *Mapt*$^{-/-}$ ○ |
| 3 | ♂ WT ● |
| 4 | ♂ WT ● |
| 5 | ♂ *Mapt*$^{-/-}$ ○ |
| 6 | ♀ WT ▲ |
| 7 | ♀ *Mapt*$^{-/-}$ △ |
| 8 | ♀ WT ▲ |
| 9 | ♀ *Mapt*$^{-/-}$ △ |
| 10 | Internal control |

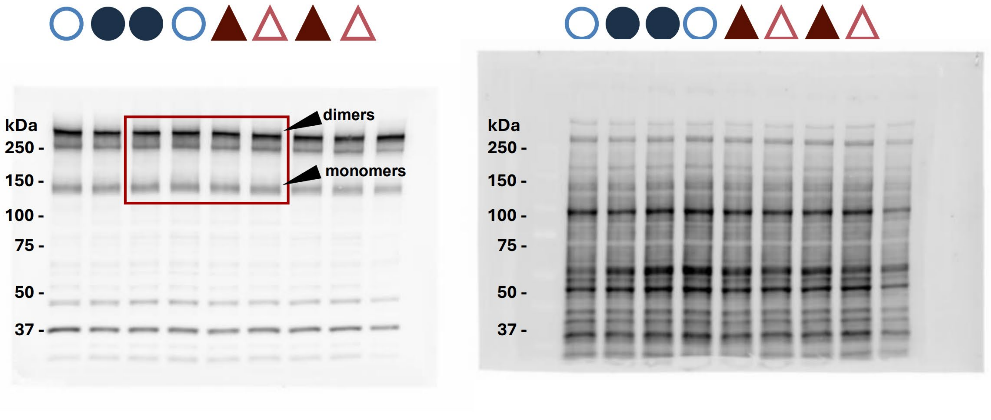


for Extended Fig. 5j-l

**s**

### Total α-tubulin (hippocampal homogenates)

| Set 2 of 4 | |
|---|---|
| Lane #: | Sample |
| 1 | Size marker |
| 2 | ♂ WT ● |
| 3 | ♂ *Mapt*$^{-/-}$ ○ |
| 4 | ♂ WT ● |
| 5 | ♂ *Mapt*$^{-/-}$ ○ |
| 6 | ♀ *Mapt*$^{-/-}$ △ |
| 7 | ♀ WT ▲ |
| 8 | ♀ *Mapt*$^{-/-}$ △ |
| 9 | ♀ WT ▲ |
| 10 | Internal control |

α-tubulin

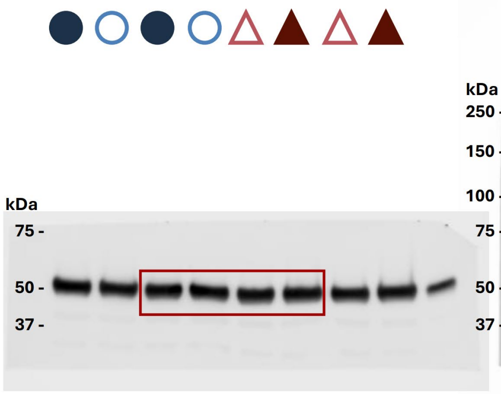


for Fig. 5a

lane total protein

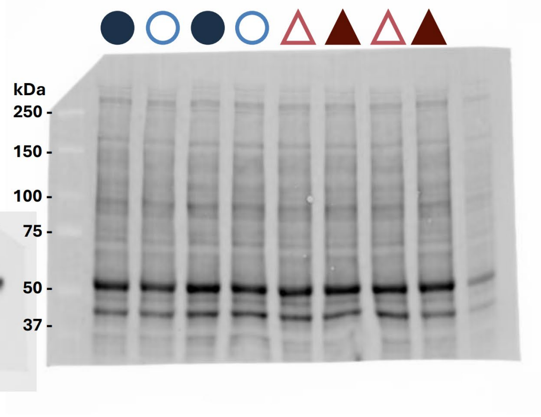


**t**

### Total β/γ-actin (hippocampal homogenates)

| Set 2 of 4 | |
|---|---|
| Lane #: | Sample |
| 1 | Size marker |
| 2 | ♂ WT ● |
| 3 | ♂ *Mapt*$^{-/-}$ ○ |
| 4 | ♂ WT ● |
| 5 | ♂ *Mapt*$^{-/-}$ ○ |
| 6 | ♀ *Mapt*$^{-/-}$ △ |
| 7 | ♀ WT ▲ |
| 8 | ♀ *Mapt*$^{-/-}$ △ |
| 9 | ♀ WT ▲ |
| 10 | Internal control |

β/γ-actin

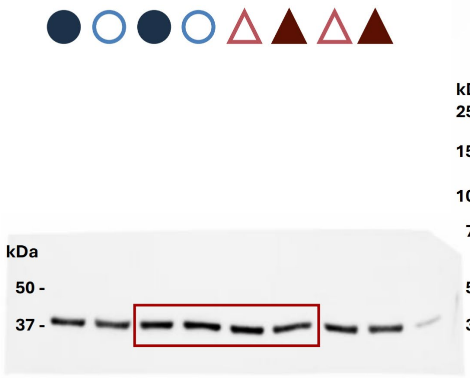


for Fig. 5b

lane total protein

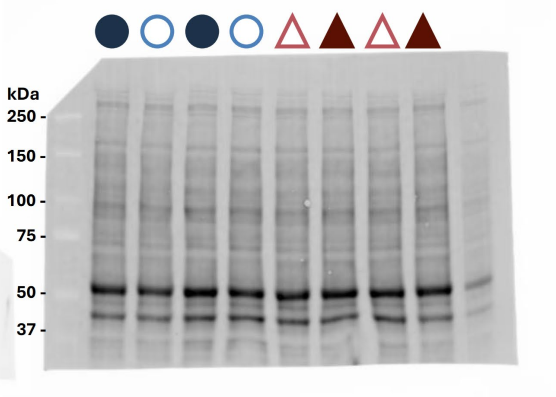


(same membrane as in panel s due to multiplexing antibodies)

**u**

### Synaptic α-tubulin (hippocampal homogenates)

| Set 2 of 4 | |
|---|---|
| Lane #: | Sample |
| 1 | Size marker |
| 2 | ♂ WT ● |
| 3 | ♂ *Mapt*$^{-/-}$ ○ |
| 4 | ♂ WT ● |
| 5 | ♂ *Mapt*$^{-/-}$ ○ |
| 6 | ♀ *Mapt*$^{-/-}$ △ |
| 7 | ♀ WT ▲ |
| 8 | ♀ *Mapt*$^{-/-}$ △ |
| 9 | ♀ WT ▲ |
| 10 | Internal control |

α-tubulin

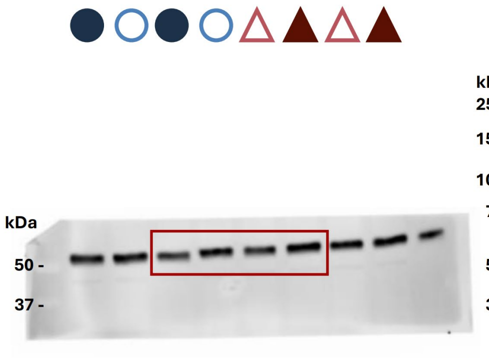


Fig. 5c

lane total protein

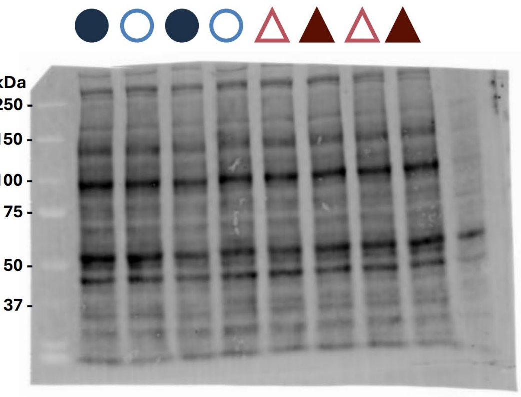

**v**

### Synaptic β/γ-actin (hippocampal homogenates)

| Set 2 of 4 | |
|---|---|
| **Lane #:** | **Sample** |
| 1 | Size marker |
| 2 | ♂ WT |
| 3 | ♂ *Mapt*$^{-/-}$ |
| 4 | ♂ WT |
| 5 | ♂ *Mapt*$^{-/-}$ |
| 6 | ♀ *Mapt*$^{-/-}$ |
| 7 | ♀ WT |
| 8 | ♀ *Mapt*$^{-/-}$ |
| 9 | ♀ WT |
| 10 | Internal control |

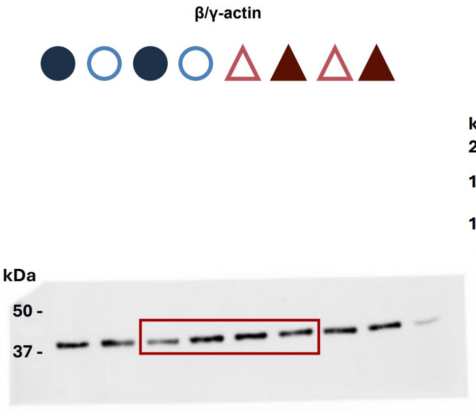


for Fig. 5d

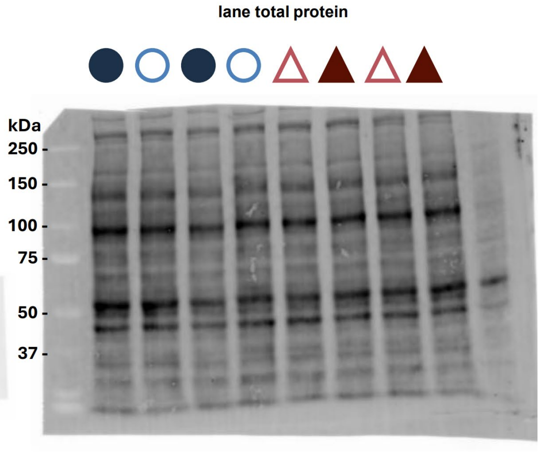


**w**

### Total tau (whole brain homogenates)

| Lane #: | Sample |
|---|---|
| 1 | Size marker |
| 2 | Size maker |
| 3 | ♂ WT |
| 4 | ♂ *Mapt*$^{+/-}$ |
| 5 | ♂ *Mapt*$^{-/-}$ |
| 6 | ♀ WT |
| 7 | ♀ *Mapt*$^{+/-}$ |
| 8 | ♀ *Mapt*$^{-/-}$ |
| 9 | Internal control |
| 10 | Size Maker |

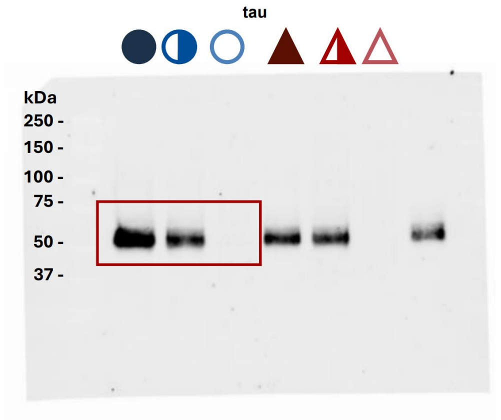


for Extended Data Fig. 1b

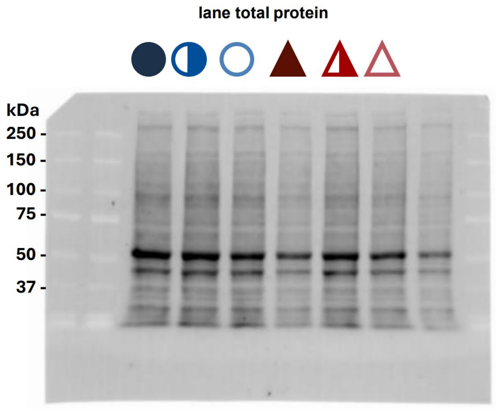

**Supplementary Methods Table 1**. Antibodies used for *Mapt*$^{-/-}$ rat immunoblotting experiments.

| Target: | Titer: | Host: | Cat #: | Supplier: |
|---|---|---|---|---|
| Tau | 1:1000 | Mouse | MABN2406 | Millipore Sigma |
| MAP1A/B | 1:500 | Rabbit | PA5-78052 | Invitrogen |
| MAP2A/B/C | 1:125 | Mouse | MA5-12826 | ThermoFisher Scientific |
| GluN2A | 1:500 | Mouse | MA527692 | ThermoFisher Scientific |
| pGluN2A Y1426 | 1:500 | Rabbit | 4206 | Cell Signaling Technology |
| GluN2B | 1:1000 | Mouse | 66565-1 | ThermoFisher Scientific |
| pGluN2B Y1472 | 1:1000 | Rabbit | 4208 | Cell Signaling Technology |
| GluA1 | 1:100 | Mouse | MAB2263 | Millipore Sigma |
| pGluA1 S845 | 1:1000 | Rabbit | AB5849 | Millipore Sigma |
| GluA2 | 1:500 | Mouse | MAB397 | Millipore Sigma |
| pGluA2 S880 | 1:1000 | Rabbit | AP0257 | ABclonal |
| mGlu1 | 1:100 | Mouse | 610965 | BD Transduction Laboratories |
| mGlu5 | 1:1000 | Rabbit | A9819 | ABclonal |
| β/γ-actin | 1:5000 | Mouse | 3700 | Cell Signaling Technology |
| α-tubulin | 1:5000 | Rabbit | 2144 | Cell Signaling Technology |
| StarBright Blue 520 Anti-Ms | 1:3000 | Goat | 12005866 | Bio-Rad |
| StarBright Blue 700 Anti-Ms | 1:3000 | Goat | 12004158 | Bio-Rad |
| StarBright Blue 520 Anti-Rb | 1:3000 | Goat | 12005869 | Bio-Rad |
| StarBright Blue 700 Anti-Rb | 1:3000 | Goat | 12004161 | Bio-Rad |
